# Complex dynamics in simplified neuronal models: reproducing Golgi cell electroresponsiveness

**DOI:** 10.1101/378315

**Authors:** Alice Geminiani, Claudia Casellato, Francesca Locatelli, Francesca Prestori, Alessandra Pedrocchi, Egidio D’Angelo

## Abstract

Brain neurons exhibit complex electroresponsive properties - including intrinsic subthreshold oscillations and pacemaking, resonance and phase-reset - which are thought to play a critical role in controlling neural network dynamics. Although these properties emerge from detailed representations of molecular-level mechanisms in “realistic” models, they cannot usually be generated by simplified neuronal models (although these may show spike-frequency adaptation and bursting). We report here that this whole set of properties can be generated by the *extended generalized leaky integrate-and-fire* (E-GLIF) neuron model. E-GLIF derives from the GLIF model family and is therefore mono-compartmental, keeps the limited computational load typical of a linear low-dimensional system, admits analytical solutions and can be tuned through gradient-descent algorithms. Importantly, E-GLIF is designed to maintain a correspondence between model parameters and neuronal membrane mechanisms through a minimum set of equations. In order to test its potential, E-GLIF was used to model a specific neuron showing rich and complex electroresponsiveness, the cerebellar Golgi cell, and was validated against experimental electrophysiological data recorded from Golgi cells in acute cerebellar slices. During simulations, E-GLIF was activated by stimulus patterns, including current steps and synaptic inputs, identical to those used for the experiments. The results demonstrate that E-GLIF can reproduce the whole set of complex neuronal dynamics typical of these neurons - including intensity-frequency curves, spike-frequency adaptation, depolarization-induced and post-inhibitory rebound bursting, spontaneous subthreshold oscillations, resonance and phase-reset, - providing a new effective tool to investigate brain dynamics in large-scale simulations.

## 1 Introduction

The causal relationship between components of the nervous system at different spatio-temporal scales, from subcellular mechanisms to behavior, still needs to be disclosed and this represents one of the main challenges of modern neuroscience. To this aim, bottom-up modeling is an advanced strategy that allows to propagate low-level cellular phenomena into large-scale brain networks (D’Angelo and Gandini Wheeler-Kingshott, 2017; Markram, 2013; Markram et al., 2015). Precise biophysical representations can be generated by “realistic” neuron and network models, but these need then to be simplified to achieve computational efficiency (D’Angelo et al., 2016a; Gerstner and Naud, 2009). Simplified neuron models are fundamental for studying the emergent properties of neural circuits in large-scale simulations and for summarizing in a principled way the electrophysiological intrinsic neural properties that drive network dynamics and high-level behaviors (Gerstner et al., 2014). The specific electroresponsive properties of single neurons are crucial for efficient signal processing, e.g. contributing to noise filtering, signal coding and synaptic plasticity. The expression of detailed neuron dynamics in simplified models would allow to analyze physiological and pathological phenomena of spiking networks during closed-loop simulations, allowing the inference of causal relationships across scales. A critical issue is therefore to obtain simplified neuronal models, that should be at the same time biologically meaningful and computationally efficient.

A compromise between accuracy and efficiency has been reached with simplified mono-compartmental neuron models (or *point neuron* models), which neglect morphology and loose some functionalities compared to detailed multi-compartmental models but gain computational efficiency. A mono-compartmental neuron model that has been widely used as the basic element of Spiking Neural Networks (SNNs) in different brain areas is the Leaky Integrate and Fire (LIF) (Lennon et al., 2014). LIFs represent neurons as first-order capacitive circuits and embed a threshold-based reset mechanism to reproduce spiking activity (Burkitt, 2006). LIFs are able to generate simple subthreshold dynamics and spike patterns but, in their original formulation, cannot reproduce smooth spike initiation zone, firing adaptation and bursting properties. Non-linear adaptive LIFs have been developed to enhance electrophysiological realism. In the Izhikevich model of cortical neurons, the dynamics of membrane potential, *V_m_*, depends on both 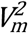 and a membrane recovery variable (Izhikevich, 2003). By introducing an exponential term in the differential equation of membrane potential and an adaptive current with slow dynamics, the action potential shape was well fitted without the need of a threshold-reset mechanism (Brette and Gerstner, 2005). However, the nonlinearity entailed more difficulties in optimizing model parameters and in computational efficiency. Therefore, recently, new linear adaptive point models have been developed (Generalized LIF, GLIF), with spike-triggered currents and moving threshold as the source of adaptation (Mihalaş and Niebur, 2009) and with stochastic processes in firing emission (Pozzorini et al., 2015; Rössert et al., 2016). The possibility to use a linear and analytically solvable neuron model is fundamental when simulating large-scale SNNs, since computational efficiency can be enhanced without severe loss in spike time accuracy and realism (Hanuschkin et al., 2010). However, GLIF can hardly generate phenomena like subthreshold oscillations, resonance and phase-reset, which are critical for large-scale network entrainment and communication (Buzsáki, 2004; Buzsáki and Draghun, 2004).

We propose here an *extended* GLIF (E-GLIF) model, which achieves a sound compromise between model complexity, biological plausibility and computational efficiency (Fig. 1). The model has a minimum number of state variables, the membrane potential and two currents, which can be associated to main biophysical subcellular mechanisms. Thanks to its mathematical structure, which is similar to GLIF and analytically solvable, E-GLIF can be optimized by traditional optimization algorithms (Pozzorini et al., 2015; Teeter et al., 2018) avoiding metaheuristic methods, like Genetic Algorithms, used for multi-compartment realistic neurons with high-dimensional parameter search space (Masoli et al., 2015). E-GLIF can reproduce a rich variety of electroresponsive properties: autorhythmicity, depolarization-induced bursting and post-inhibitory rebound bursting, specific input-output (*f-I_stim_*) relationships, spike-frequency adaptation, phase-reset, sub-threshold oscillations and resonance. A comprehensive example of this entire set of excitable properties was given by the E-GLIF of a cerebellar Golgi cell (GoC), whose electrophysiological properties have been carefully investigated and modeled previously using realistic multi-compartmental approach (Forti et al., 2006; Solinas et al., 2007a, 2007b). And since the GoC expresses among the most common and varied electroresponsive properties, the E-GLIF model should be easily applied to various central neurons that can be represented with mono-compartmental models and promote the investigation of complex brain dynamics in large-scale simulations (Jordan et al., 2018).

**Figure 1.**
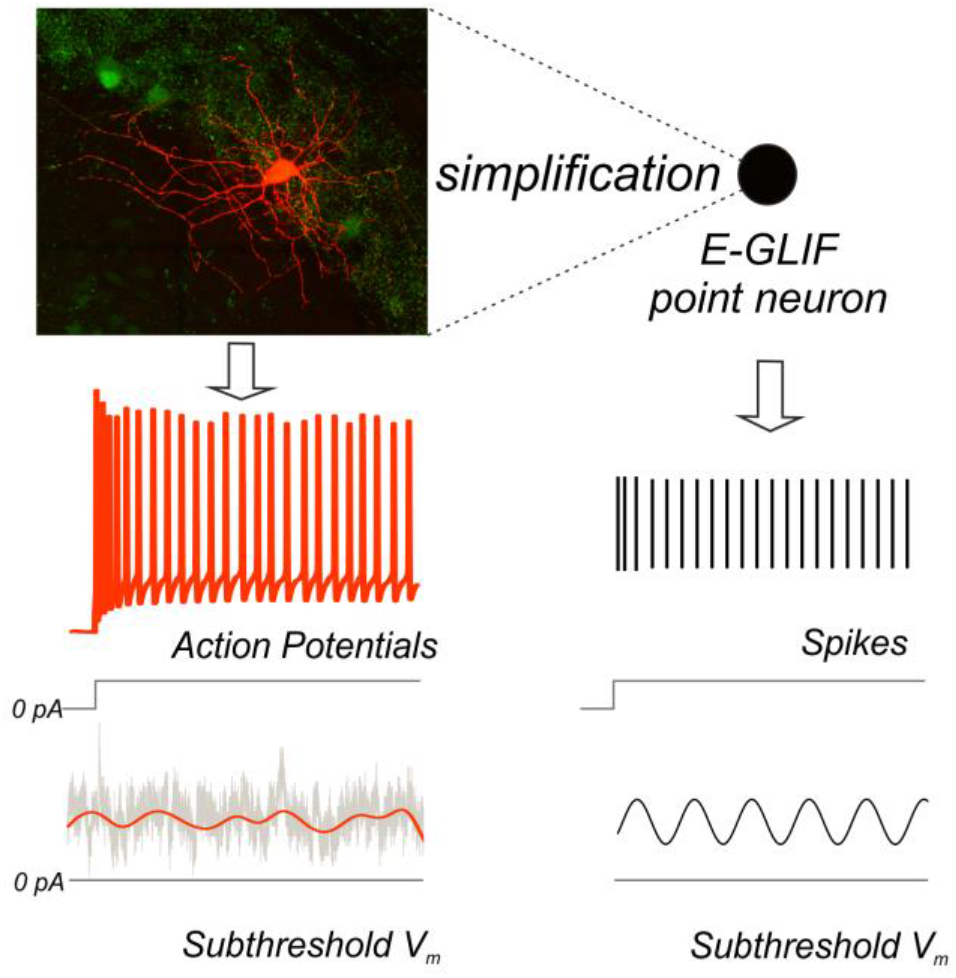
From complex to simplified neuron model. Neurons exhibit complex properties that affect synaptic plasticity, circuit dynamics and behavior. The point neuron described here shows that it is possible to capture this complexity through a simplified model. The top left panel shows a Golgi cell of mouse cerebellum stained with red tomato and reconstructed using a fluorescence confocal microscope (courtesy of Prof. Javier De Felipe, Department of Neuroanatomy and Cell Biology, Instituto Cajal (CSIC), Madrid, Spain). The Golgi cell shows a complex set of dendritic and axonal ramifications. In whole-cell patch-clamp electrophysiological recordings from the soma, the Golgi cell behaves as a pacemaker generating adapting spike trains in response to step current injection (bottom left) and other more complex properties that are explained in this paper. As shown in the picture, membrane potential firing and subthreshold properties recorded *in vitro* (left panel) can be observed in simulated spiking patterns and subthreshold dynamics obtained through the E-GLIF point neuron (right).

## 2 Methods

In this work, taking the move from previous GLIF neurons (Hertäg et al., 2012; Mihalaş and Niebur, 2009; Pozzorini et al., 2015), we have developed and tested the E-GLIF neuron. The model was implemented in the Neural Simulation Tool (NEST) (Diesmann and Gewaltig, 2002), using NESTML (Plotnikov et al., 2016) and the C++ core of NEST. Experimental recordings were performed from cerebellar GoCs using patch-clamp recording techniques for validation.

### 2.1 The model

E-GLIF couples time-dependent and event-driven algorithmic components to generate a rich set of electrophysiological behaviors, while keeping the advantages of LIF neuron models in terms of simplicity and analytical solvability. The E-GLIF neuron includes 3 linear Ordinary Differential Equations (ODEs) describing the time evolution of membrane potential (*V_m_*) and of two intrinsic currents (*I_adapt_* and *I_dep_*). Each of these three state variables is modified by an update rule at spike events, which are generated according to a probabilistic threshold crossing controlled by an escape noise.

The model is defined as follows:

- State variables evolution (second-order 3-dimenstional ODE system):

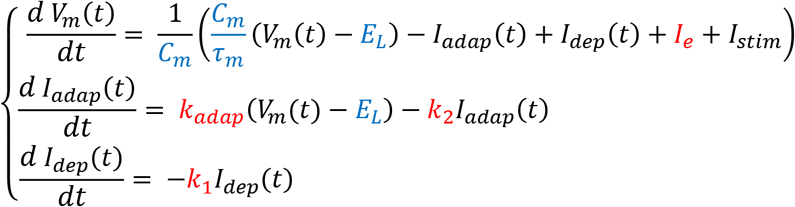 Where:

*I_stim_* = external stimulation current;
*C_m_* = membrane capacitance;
*τ_m_* = membrane time constant;
*E_L_* = resting potential;
*I_e_* = endogenous current;
*k_adap_*, *k*_2_ = adaptation constants;
*k*_1_ = *I_dep_* decay rate.
- Spike generation: the spike generation rule is stochastic, depending on the escape rate function *λ*(*t*) (Gerstner and Kistler, 2002; Jolivet et al., 2006) and the refractory interval Δ*t_ref_*:

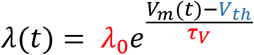

where:

*V_th_* = threshold potential;
*λ*_0_, *τ_V_* = escape rate parameters. If

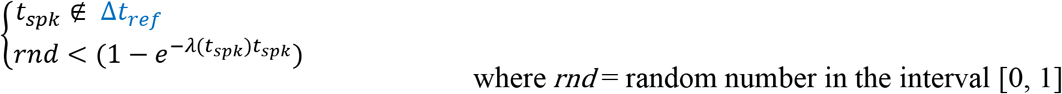

then, a spike is generated at time *t_spk_*, according to a point process. Thus, spike times do not correspond strictly to when the potential reaches the threshold: they occur with higher probability if the membrane potential is near the threshold, depending on the parameters *τ_V_* and *λ_0_* that define the minimum distance from threshold corresponding to the maximum probability of having a spike (Gerstner et al., 2014).
- Update: following a spike, the state variables are modified according to the rules:

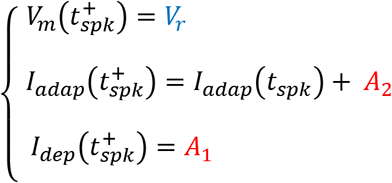

where:

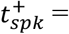 time instant immediately following the spike time *t_spk_*
*V_r_* = reset potential;
*A*_2_, *A*_1_ = model currents update constants.

The parameters in the model include those directly related to neurophysiological quantities (*C_m_*, *τ_m_*, *E_L_*, *Δt_ref_*, *V_th_*, *V_r_* in blue), that are fixed for each specific cell type, and the more abstract ones related to neuron-specific functional mechanisms, that need to be optimized (*k_adap_, k_2_, k_1_, A_2_, A_1_, I_e_*, in red).

In addition to the leaky current term, 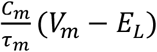 each one of the membrane currents defined in the model (*I_e_*, *I_adapt_*, *I_dep_*) accounts for a different mechanism that can be properly parameterized:

*I_e_* is an endogenous current modeling the net contribution of depolarizing ionic currents generating autorhythmicity (Mihalaş and Niebur, 2009).
*I_adap_* is an adaptive current, usually hyperpolarizing, which is characterized by a small spike-triggered increment (*A*_2_) that decays thereafter according to *k_adap_* and *k*_2_. *I_adap_* models the activation of potassium channels generating a slow hyperpolarizing current. Since *I_adap_* activates slowly while *I_dep_* is already decaying, the balance between the two currents generates spike-frequency adaptation and afterhyperpolarization. Moreover, by being coupled with *V_m_* by *k_adap_*, *I_adap_* endows the model with the capability of generating post-inhibitory rebound bursting, intrinsic subthreshold oscillations and resonance (Brette and Gerstner, 2005; Hertäg et al., 2012).
*I_dep_* is a depolarizing spike-triggered current, which has a larger spike-triggered increment (*A_1_*) and faster decay (*k_1_*) compared to *I_adap_. I_dep_* mimics the fast (almost instantaneous) activation and deactivation of sodium channels. *I_dep_* can generate depolarization-induced bursting and sustain post-inhibitory rebound bursts (Mihalaş and Niebur, 2009).

Considering the matrix form of the ODE system describing the model dynamics:

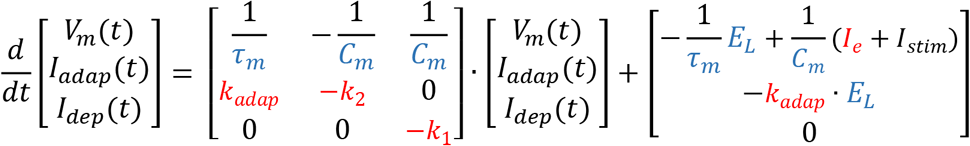

the general solution is:

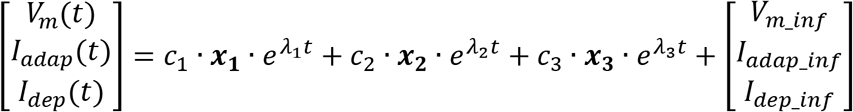

where:
*c*_1_, *c*_2_, *c*_3_ are arbitrary constants depending on the initial conditions
*λ*_1_, *λ*_2_, *λ*_3_ are the eigenvalues of the coefficient matrix, with values:

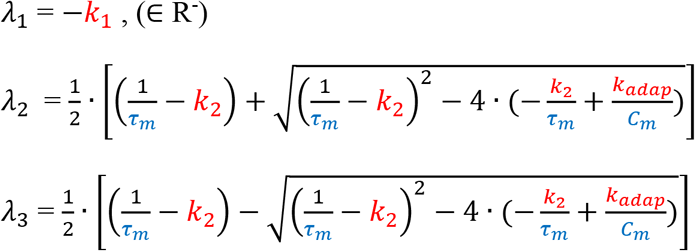

***x***_**1**_, ***x***_**2**_, ***x***_**3**_ are the eigenvectors associated to each eigenvalue *V_m_inf_, I_adap_inf_*, *I_dep_inf_* are the stationary solutions for each state variable.

For the membrane potential, the solution is (Hertäg et al., 2012):

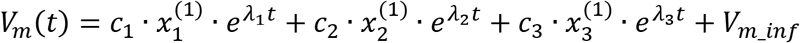

Specifically:

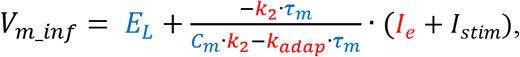

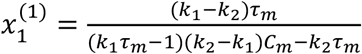, the first component of the eigenvector associated to the eigenvalue *λ*_1_.

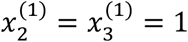, the first component of the eigenvectors associated to the eigenvalues *λ*_2_ and *λ*_3_, respectively

Considering that *k_1_* is real and positive, the dynamics of the solution depends on the discriminant:

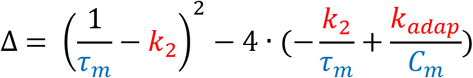

<u>Case 1:</u>exponential and stable solution; the following conditions need to be verified:

- Δ > 0 ⇒ *λ*_1_, *λ*_2_, *λ*_3_ ∈ R (the solution is exponential)
- *λ*_1_, *λ*_2_, *λ*_3_ < 0 (the solution is stable)

In addition, we need to verify that *V_m_inf_* ∝ *I_e_*+*I_stim_*, i.e. *V_m_inf_* is proportional to the total input current through a positive coefficient, to have a coherent value of steady state membrane potential.

These conditions result in the following constraints on parameters:

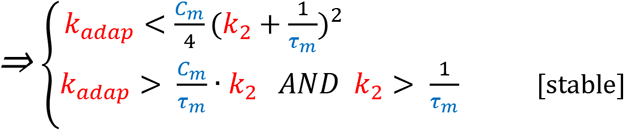

<u>Case 2:</u> oscillatory solution; the following conditions need to be verified:

- Δ < 0 ⇒ *λ*_1_, *λ*_2_, *λ*_3_ *∈ C* (the solution is oscillatory);
- If *Re[λ*_1_, *λ*_2_, *λ*_3_*] = 0* ⇒ the oscillations have null damping;
- If *Re[λ*_1_, *λ*_2_, *λ*_3_*] < 0* ⇒ the oscillations are damped and the solution is stable.

Analogously to the previous case, we need to verify that *V_m_inf_* ∝ *I_e_*+*I_stim_* through a positive coefficient.

The resulting constraints among parameters are:

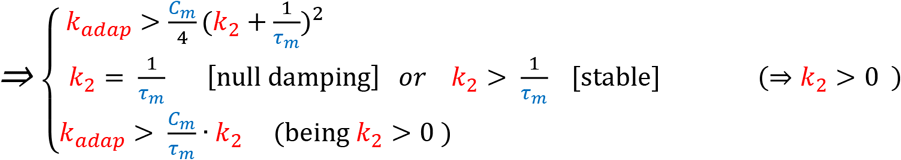

In this case, oscillations depend on the imaginary part of the eigenvalues 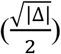 and thus have angular frequency 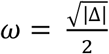 and related frequency 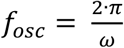.

As reported in Figure 2A, these parameter constraints define two regions in the *k_2_-k_adap_* plane, corresponding to exponential and stable solutions (green area in the figure) and oscillatory not damped solutions (red line in the figure) for self-sustained oscillations of the membrane potential.

**Figure 2.**
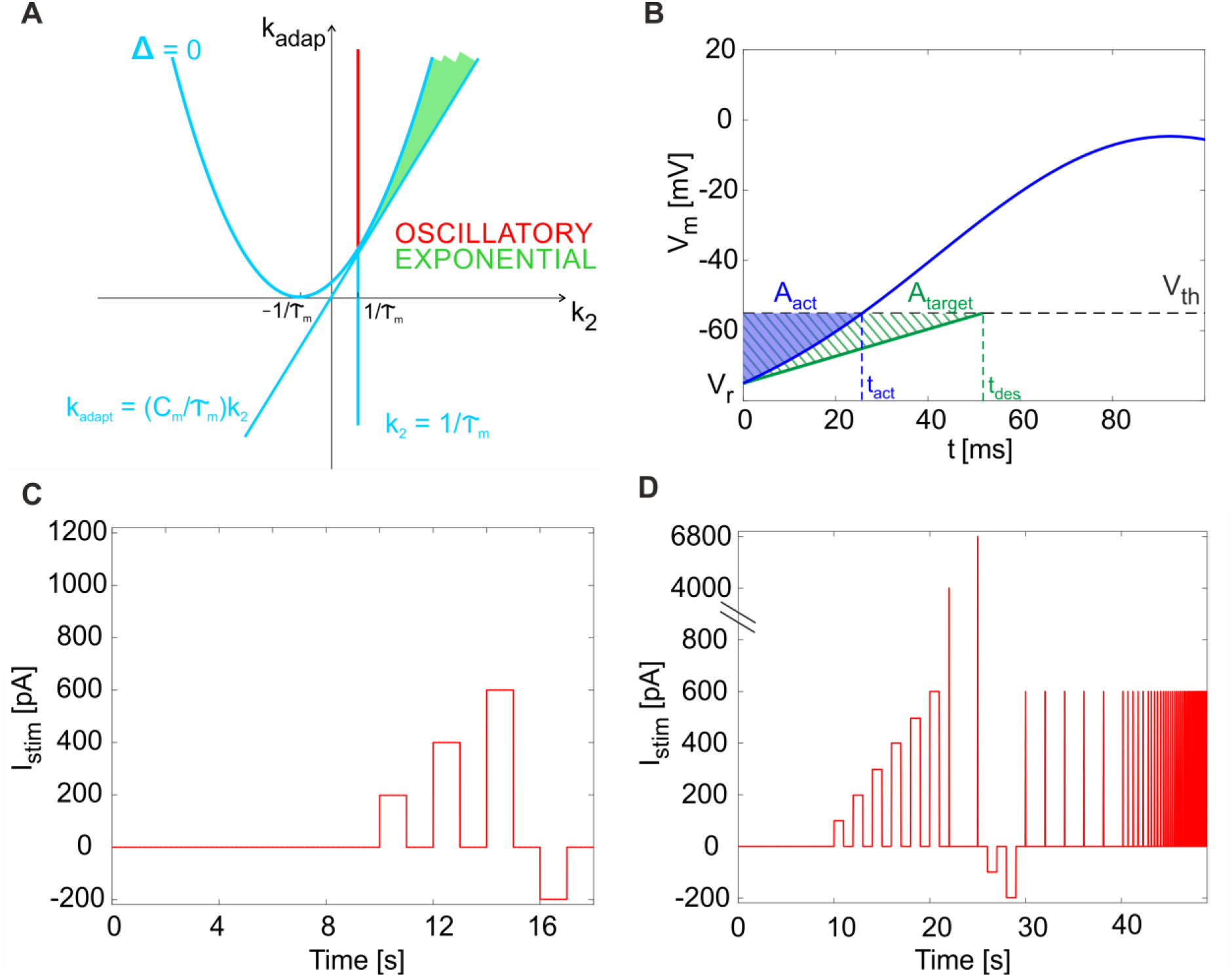
Mathematical properties of the model. (A) Parameter plane. The solution types as a function of parameters *k_adap_* and *k_2_*. The light blue lines identify the critical values separating sub-planes where the solution type changes. Green area: exponential and stable solutions (where Δ > 0 and *k_adap_* > (C_m_/τ_m_)·*k*_2_ and *k*_2_ > 1/τ_m_). Red segment: oscillatory and not damped solutions (where Δ < 0 and *k*_2_ = 1/*τ_m_*). (B) Cost function evaluation exemplified by showing *V_m_*(*t*) within one time interval (from relative initial time t_0_ = 0). Blue: the actual *V_m_*(*t*), crossing the threshold (*V_th_*) at spike time *t_act_*. Green dashed: the simplified target *V_m_*(*t*), crossing the threshold (*V_th_*) at spike time *t_des_*. Both *V_m_*(*t*) curves start from reset value (*V_r_*). The error is computed as the difference of blue and green areas, neglecting trend above *V_th_*. (C) The short continuous stimulation protocol is represented. This is used for assessing model optimization, applying the same input current values used during optimization, i.e. a zero-current phase, 3 depolarizing steps with input *exc1, exc2, exc3* = 200 pA, 400 pA, 600 pA and a hyperpolarizing step with input *inh* = −200 pA, all interleaved with zero-current phases. (D) The full multi-phase *in vitro* stimulation protocol is reported. This has been used for the continuous long simulations (48.93 seconds) to validate the model and consequently for the input definition in the experimental protocol. After 10 s at zero current (spontaneous activity), 6 increasing current steps are imposed (from 100 to 600 pA, with 100 pA of increase), each lasting one second; they are interleaved by 1 s of zero-current. Then, two impulses are provided (4000 pA and 6800 pA, each lasting 0.5 ms). Then, 2 negative current steps are imposed (−100 pA and −200 pA), each lasting 1 s. Finally, 13 sequences of pulse trains are provided. Each sequence is made up of 5 pulses (each 600 pA lasting 30 ms); the time interval between pulses is constant within each sequence (2, 0.5, 0.28, 0.2, 0.17, 0.13, 0.1, 0.08, 0.06 ms).

### 2.2 Optimization

#### 2.2.1 Model parameters

To generate a neuron-specific model, we first considered the parameters that are directly measured as neurophysiological quantities (*fixed parameters*, highlighted in blue in Section 2.1): *C_m_, τ_m_, E_L_, Δt_ref_, V_th_, V_r_* were fixed to biological values taken from literature or available from animal experiments or databases (Tripathy et al., 2014). For the other neuron-specific functional parameters (*tunable parameters*, highlighted in red in Section 2.1), *k_adap_, k_2_, k_1_, A_2_, A_1_, I_e_*, we developed an optimization strategy based on a desired input-output relationship, considering a current step *I_stim_* as the input and spike times as the output. Specifically, we supposed to evaluate the neuron response to the inputs listed in Table 1:

- *I_stim_* = 0: zero current (*zero_stim*) generating spikes at frequency *tonic_freq* to evaluate autorhythm;
- *I_stim_* at three increasing excitatory current steps (*exc1* < *exc2* < *exc3*) producing firing with increasing frequency (*freq1* < *freq2* < *freq3*) to reproduce the *f-I_stim_* relationship and spike- frequency adaption (i.e. steady-state decreased frequency with *gain1* > *gain2* > *gain3*);
- *I_stim_* = *inh*, an inhibitory input current to evaluate the occurrence of an inhibition-induced silence followed by rebound burst, made of at least 2-spikes.

**Table 1.**
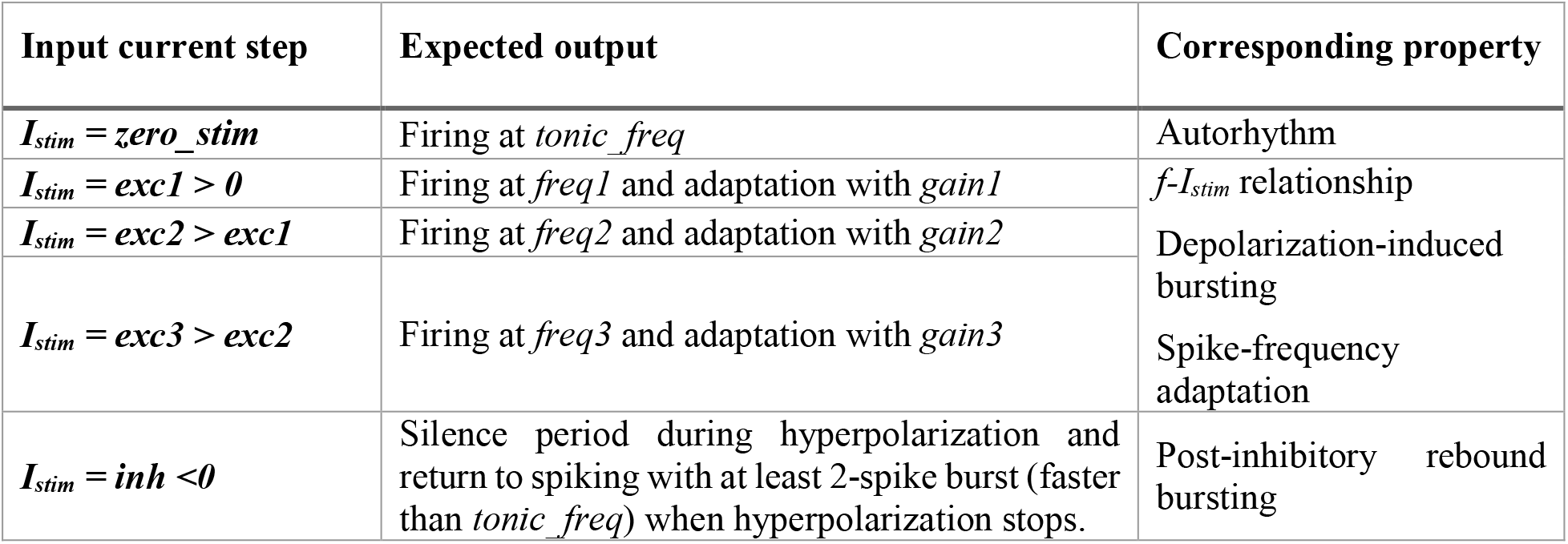
Inputs and expected outputs for the optimization algorithm with corresponding target electrophysiological property.

#### 2.2.2 Cost function, constraints and algorithm

To evaluate different parameter sets by computing the corresponding cost function, we exploited the analytical tractability of the model and we evaluated the model solution *V_m_(t)*, within the most significant time windows (initial, transitory, and steady state) during each stimulation current step. For each *I_stim_* = (*i*) = *zero_stim, exc1, exc2, exc3*, the three time windows taken into account were: 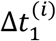, from 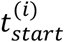 to first spike 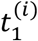, 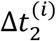 between 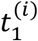 and second spike time 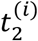, and 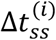 between two spikes at steady-state.

At the beginning of each depolarizing phase (i.e. during 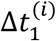), we supposed the state variables to be initialized as follows:

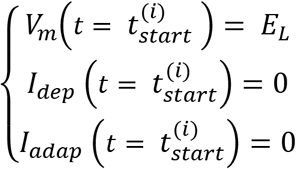

Then, for the following time intervals, the initial conditions were derived from the update rules of the model, supposing that the system had reached the steady-state condition when the adaptive current, *I_adap_*, decayed during the inter-spike interval of an amount equal to *A_2_*, i.e. its update constant.

Analogously, to evaluate rebound bursting, we computed the solutions with *I_stim_* = 0, just after a hyperpolarization (*I_stim_* = *inh* < 0), within 2 consecutive time windows: 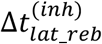, from the end of the inhibitory current stimulus to the first rebound spike time *t_lat_reb_*, and during the 1^st^ Inter-Spike Interval (ISI) of rebound burst 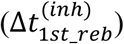. Again, the initial conditions were derived using the update rules.

Starting from all the solutions computed in terms of *V_m_*(*t*), for each input *I_stim_* in Table 1, we derived the desired spike times 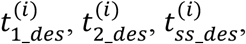, or 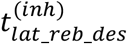 and 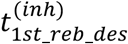 by imposing a spike event at the time when *V_m_*(*t*) = *V_th_*. Therefore, the spike generation during optimization was assumed deterministic. To take into account the variability in spike generation due to the stochasticity into the model, for each stimulation input during training, we used a distribution of 10 desired firing frequencies with specific mean and Standard Deviation (SD), and thus a distribution instead of a single target spike time.

The parameters *k_adap_, k_2_, A_2_, k_1_, A_1_, I_e_* were optimized through the Sequential Quadratic Programming (SQP) algorithm from the MATLAB R2015b Optimization Toolbox, with normalized parameter values and constraints. SQP optimization aims at the simultaneous minimization of a cost and constraint function, using a gradient-based minimization method. We set the stopping criteria to the iteration when the variations in parameter search, cost or constraint functions were below 10^-3^ or the cost function reached a value lower than 0.1. We chose the cost function for gradient minimization, summing all the errors relative to the analyzed properties along the stimulation protocol as in Table 1 (zero input, 3 excitatory currents, first and second spikes of the post-inhibitory rebound burst).

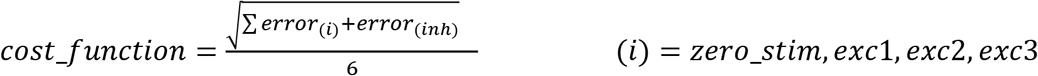

Each error term compared the actual area *A_act_*, i.e. the area between the actual *V_m_*(*t*) curve and the *V_m_*(*t*) = *V_th_* horizontal line, during each desired ISI, to the target area *A_target_* (Fig. 2B):

> 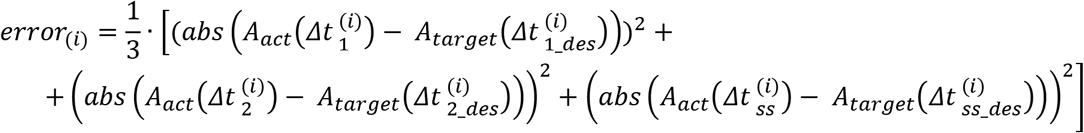
>
>
> for each input current (*i*)
> where:
>
>
> 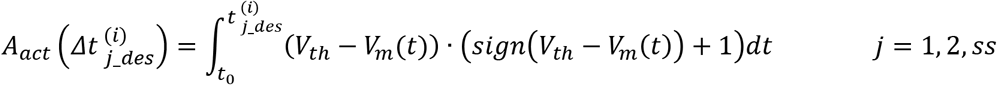
>
>
> computing only the area from the relative initial time t_0_ = 0 to the spiking time, before reaching the threshold, since spike-reset mechanisms were enabled in the model simulations.
>
> 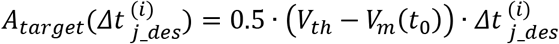
>
>
> that can be considered an approximation of the ideal sub-threshold membrane potential for a linear LIF neuron not firing at very low spiking frequencies.
>
> 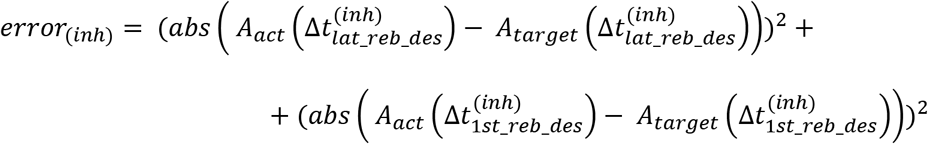
>
>
> considering two aspects of the rebound burst following inhibition, the time of the first spike after hyperpolarization and the distance of the second spike after it.

Based on the mathematical considerations in Section 2.1 about the dynamics of the solutions, the parameter space needed to be constrained to obtain the desired membrane voltage evolution (Fig. 2A). Further limits in the parameter space could be included to take into account neuron-specific information from neurophysiology.

In order to evaluate the convergence and stability of the optimization process, for each optimization run, parameters were initialized to random values within their ranges and 5 optimizations were performed with different random initializations. The median of all the resulting parameters was then considered as the final optimal set and used to run complete simulations using the simulator PyNEST (Eppler et al., 2009) for assessing the optimization results and for further validation.

### 2.3 E-GLIF model of the cerebellar Golgi cell

E-GLIF model and optimization were applied to reproduce the complex electroresponsiveness of cerebellar Golgi cells (GoC). GoCs are the main inhibitory neurons in the granular layer of cerebellum and are responsible for reshaping the input signals coming from mossy fibers. In single-cell recordings, GoCs show spontaneous firing around 8 Hz, a nearly-linear input-output relationship (about 0.25 Hz/pA), input-dependent spike-frequency adaptation when depolarization is maintained, rebound bursting after hyperpolarization, phase-resetting, subthreshold self-sustained oscillations and resonance in the theta band (around 3-6 Hz) (D’Angelo et al., 2016a; Forti et al., 2006; Solinas et al., 2007a, 2007b). A multi-compartmental realistic model (Solinas et al., 2007a, 2007b) assumed that dendrites were passive and used them to redistribute the passive electrotonic load while placing all the ionic channels in the soma, suggesting that an appropriate single point model could have been effective as well. In the present E-GLIF model, all electrical properties are collapsed in a point and gating kinetics of ionic channels are substituted by lumped and simplified membrane mechanisms.

#### 2.3.1 Model construction and optimization: physiological parameters, cost function and constraints

As reported in Section 2.2.1, the values of the electrophysiological parameters were taken from literature (Forti et al., 2006; Solinas et al., 2007a, 2007b) and databases (Tripathy et al., 2014) describing experimental GoC properties *in vitro*; their values are listed in Table 2.

**Table 2.** Electrophysiological parameters of the cerebellar GoC taken from literature.

| Parameter | Value | Reference |
| --- | --- | --- |
| $C_m$ | 145 pF | (Solinas et al., 2007a) |
| $\tau_m (= C_m * R_{in})$ | 44 ms | (Solinas et al., 2007a) |
| $E_L$ | -62 mV | (Solinas et al., 2007a) |
| $\Delta t_{ref}$ (spike width) | 2 ms | (Tripathy et al., 2014) |
| $V_r$ | -75 mV | (Tripathy et al., 2014) |
| $V_{th}$ | -55 mV | (Forti et al., 2006) |

The optimization of the remaining tunable parameters was achieved setting proper values to the target behaviour of the optimization algorithm, derived from literature (Forti et al., 2006; Solinas et al., 2007a, 2007b), as shown in Table 3.

**Table 3.**
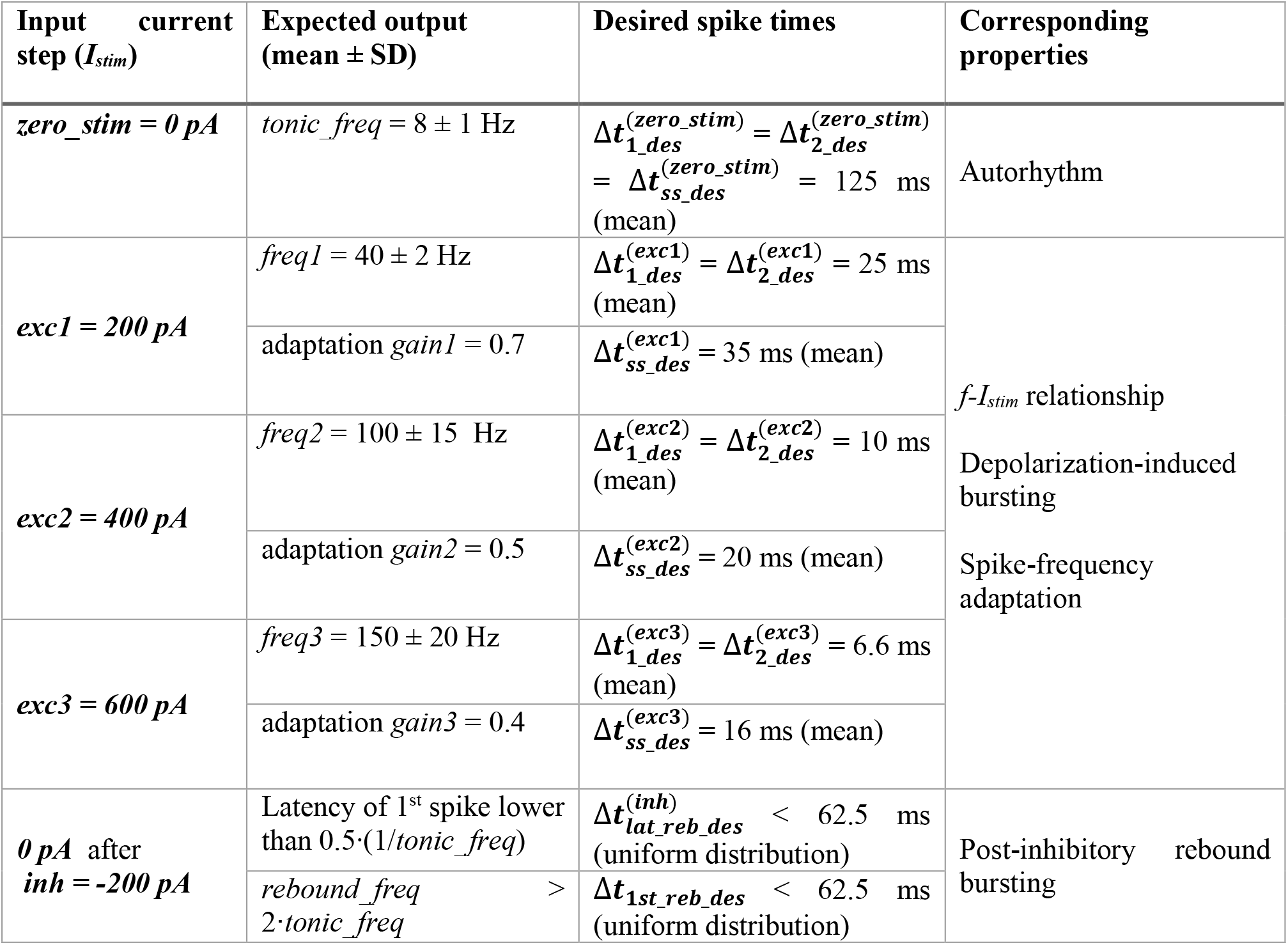
Input-output relationship for GoC model optimization with corresponding electroresponsive properties (Forti et al., 2006; Solinas et al., 2007a, 2007b). Input: current step values (*zero_stim, exc1, exc2, exc3, inh*). Output: the distribution of desired firing rates during the autorhythmic phase (*tonic_freq*), at the beginning and end of depolarizing phases (*freq1, freq2, freq3*), and during the rebound burst (latency and *rebound_freq* compared to *tonic_freq*). For each output, the desired spike intervals in the significant time windows 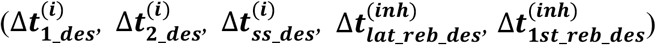 are computed for each input current (superscript in brackets).

In order to account for the whole-set of GoC electrophysiological properties, the cost function included all the terms reported in *cost_function* in Section 2.2.2. In addition, the parameter space was limited to fulfil mathematical and neurophysiological constraints:

Nonlinear constraints:

> Negative discriminant (Δ < 0) to obtain an oscillatory membrane potential:
>
>
> 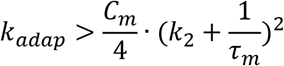
>
> Controlled *V_m_* oscillation frequency: 3 < *fosc* < 8 *Hz*, where *fosc* is defined in Section 2.1.
>
> Controlled amplitude of oscillations (|*V_m_osc_*|), during the intervals 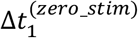, 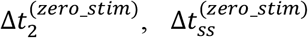 of the zero-current phase (*I_stim_* = 0) and during the hyperpolarizing interval *hyp* (with *I_stim_* = *inh*):
>
>
> 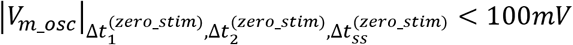
>
>
>
>
> 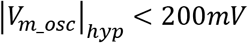
>
>
> 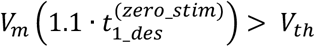 to constraint the occurrence of the first spike event during the zero-current phase, so to trigger all the spike-reset-update mechanisms.

Linear constraints:

> GoC show faster dynamics of the sodium ion current with respect to potassium one (accounted for by the current updates *A_1_* and *A_2_*):
>
>
> 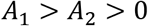

Parameter bounds:

> Not-damped *V_m_* oscillations:
>
>
> 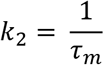
>
> After Δ*t_ref_*, negligible contribution of the depolarizing current *I_dep_*, during the zero-current phase (*I_stim_* = 0), to avoid interference with the neuron spontaneous activity, being *I_dep_* the depolarizing spike-triggered current, with decay *k_1_*:
>
>
> 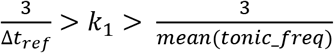
>
> Realistic values of *I_dep_* and *I_adap_* based on the neurophysiological values of sodium and potassium ion currents (Solinas et al., 2007a):
>
>
> 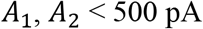
>
> Limited values of the endogenous current:
>
>
> 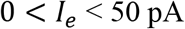

The optimal parameter set was chosen as the median of the final values over the 5 optimization runs with different random parameter initialization.

#### 2.3.2 Model simulations

Following parameter optimization in MATLAB, we derived irregular firing parameters (*λ_0_* and *τ_V_*), with the aim to obtain a physiological variability of spike events during autorhythmicity. In fact, in the PyNEST model simulations, we activated the firing stochasticity that was disabled during optimization. Then, the GoC simulations in NEST proceeded in two phases.

First, we implemented a multi-step protocol with the same input currents used for optimization, taken from literature (see Table 3 and Par. 2.3.1). This phase was fundamental to assess the effectiveness of parameter tuning in continuous simulations with escape noise during application of dynamic input patterns, and not only in sample intervals (i.e. 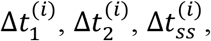 or 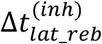 and 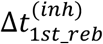). In these sequences, after a 10-sec zero-current phase used to evaluate firing irregularity, we delivered three 1-sec steps of increasing amplitude (*exc1, exc2, exc3* = 200, 400, 600 pA) to monitor the *f-I_stim_* slope and spike-frequency adaptation, interleaved with 1-sec zero-current to let the neuron recover to its spontaneous activity. We then stimulated the neuron with a 1-sec inhibitory step current (−200 pA) for evaluating rebound bursting during a subsequent zero-current step (Fig. 2C).

Secondly, we added stimulation patterns required to evaluate the emergence of model features (autorhythmicity, rebound bursting, *f-I_stim_*, adaptation) at higher resolution and to check for further emergent properties that were not considered during the optimization process (e.g. resonance, phase-reset), including:

- an initial zero-current phase of 10 s to evaluate frequency and irregularity of intrinsic firing;
- six depolarizing steps lasting 1 s, with input currents ranging from 100 pA to 600 pA (increments of 100 pA), interleaved with 1-sec zero-current phases, to test intrinsic excitability;
- two zero-input phases lasting 2.5 s, where we provided a short pulse excitation (amplitude 4 nA and 6.8 nA, for 0.5 ms), to measure phase-reset mechanism;
- two 1-sec hyperpolarizing phases with inhibitory current −200 pA and −100 pA, followed by a 1-sec zero-stimulation phase, where evaluating rebound bursting properties;
- a sequence of 5 steps, each of amplitude 600 pA, lasting 30 ms, at increasing frequencies: 0.5-2-3.5-5-6.3-7.7-10-12-15 Hz, to evaluate resonance.

The resulting total duration of the whole protocol was 48.93 seconds (Fig. 2D). Each simulation with the optimized Golgi neuron was run 10 times with different seeds of the random number generator used to produce the escape noise, and thus spike stochasticity.

#### 2.3.3 Model synaptic activation

Synaptic mechanisms were added to the model in order to allow its connection with different input neural populations and simulate the GoC response to network activity. The synapses were conductance-based, with an input spike-triggered conductance change according to an alpha function (Roth and van Rossum, 2013). This kind of synaptic model was chosen to maximize the realism in synapses behavior, despite losing the neuron model linearity when connecting to other neurons and increasing the computational load. This solution can be considered acceptable when using a small-/medium-scale SNN. However, the NEST platform flexibility guarantees the possibility to use the E-GLIF with current-based synapses, which are less realistic but also require a lower computational load and can be suitable for large-scale SNN simulations (Cavallari et al., 2014). For synaptic testing, GoC E-GLIF was connected to two different spike generators, excitatory and inhibitory, respectively. We provided a 50 Hz spike train on the excitatory synapse lasting for 800 ms, while the inhibitory synapse received a short spiking burst, for evaluating the capability of the neuron model to produce a rebound burst after an inhibitory spiking input.

#### 2.3.4 Experimental data acquisition and analysis

The outcome of simulations using the long multi-phase protocol was compared to real data acquired *ad hoc* from mice GoCs in acute cerebellar slices. These data are not identical to those derived from literature, allowing therefore to test the generalization capability of GoC E-GLIF beyond the specific dataset used for model construction. As for the model, also for the data we evaluated the same electrophysiological properties and used similar inputs to the neurons described in Section 2.3.2 (see the protocol in Figure 2D).

The experiments have been conducted on 16-to-21-day-old (P0=day of birth) male and female mice heterozygous for the bacterial artificial chromosome insertion of EGFP under the control of the glycine transporter type 2 gene (Zeilhofer et al., 2005) (GlyT2-GFP mice). All procedures were conducted in accordance with European guidelines for the care and use of laboratory animals (Council Directive 2010/63/EU), and approved by the ethical committee of Italian Ministry of Health (628/2017-PR). The mice were anesthetized with halothane (1 ml in 2 L administered for 1–2 min) and killed by decapitation in order to remove the cerebellum for acute slice preparation according to a well-established technique (Cesana et al., 2013; Forti et al., 2006). The cerebellum was gently removed, and the vermis was isolated and fixed on the stage of a vibroslicer (Leica VT1200S) with cyanoacrylic glue. Acute 220-μm-thick slices were cut in the parasagittal plane in cold Kreb’s solution and maintained at 32 °C before being transferred to a 1.5 ml recording chamber mounted on the stage of un upright epifluorescence microscope (Axioskop 2 FS; Carl Zeiss, Oberkochen, Germany) equipped with a 63, 0.9 NA water-immersion objective. Slices were perfused with Kreb’s solution and maintained at 32°C with a Peltier feedback device (TC-324B, Warner Instrument Corp., Hamden, CT). The Kreb’s solution contained the following (in mM): 120 NaCl, 2 KCl, 1.2 MgSO_4_, 26 NaHCO3, 1.2 KH_2_PO_4_, 2 CaCl_2_, and 11 glucose, and was equilibrated with 95% O_2_ and 5% CO_2_, for pH 7.4.

Whole-cell patch-clamp recordings were performed with Multiclamp 700B (−3dB; cutoff frequency 10 kHz), sampled with Digidata 1550 interface, and analyzed off-line with pClamp10 software (Molecular Devices). Patch pipettes were pulled from borosilicate glass capillaries (Sutter Instruments, Novato, CA) and filled with internal solution containing (in mM): potassium gluconate 145, KCl 5, HEPES 10, EGTA 0.2, MgCl_2_ 4.6, ATP-Na_2_ 4, GTP-Na_2_ 0.4, adjusted at pH 7.3 with KOH. Pipettes had a resistance of 3–5 MΩ when immersed in the bath. Signals were low-pass filtered at 10 kHz and acquired at 50 kHz. The stability of whole-cell recordings can be influenced by modification of series resistance (R_s_). To ensure that R_s_ remained stable during recordings, passive electrode-cell parameters were monitored throughout the experiments.

In each recording, once in the whole-cell configuration, the current transients elicited by 10 mV hyperpolarizing pulses from the holding potential of −70 mV in voltage-clamp mode showed a biexponential relaxation. The recording properties for 5 cells from 3 mice were measured as follows (values reported as mean ± Standard Error of Mean): membrane capacitance was evaluated from the capacitive charge (37.1 ± 8.5 pF), while the membrane resistance was computed from the steady-state current flowing after termination of the transient (195.9 ± 97.8 GΩ). The 3 dB cutoff frequency of the electrode-cell system, f_VC_, was calculated as f_VC_ = (2*π* • 2τ_VC_)^-1^, with τ_VC_ = 215.7 ± 7.4 μs, resulting in 0.7 ± 0.02 kHz. These properties were constantly monitored during recordings. Cells showing variation of R_s_ higher than 20% were discarded from analysis.

After switching to current clamp, GoCs were maintained at resting membrane potential by setting the holding current at 0 pA. Intrinsic excitability was investigated by injecting 2-sec steps of current (from −200 to 600 pA in a 100-pA increment). Resonance was investigated by applying sequences of 30-ms 600 pA current steps repeated 5 times at different frequencies (range from 2 Hz to 10 Hz) or injecting a single 0.5-ms step of 4 nA. Finally, the cells were maintained at their resting membrane potential for evaluating subthreshold *V_m_* oscillations.

Using an automatic spike detection algorithm, spike times were extracted from the experimental recordings, and used for electrophysiological feature extraction (see Section 2.3.5).

#### 2.3.5 Feature extraction

To evaluate GoC electroresponsive behavior, multiple parameters were computed according to (Solinas et al., 2007a, 2007b), from both simulation and recording spike trains:

- the tonic firing rate, *f_tonic_*, was the inverse of the mean ISI, during the initial zero-current phase;
- the coefficient of variation of inter-spike intervals (CV_ISI_) was measured during the 10-sec zero-input step to quantify the irregularity of firing;
- the firing rate, *f*, was the inverse of the mean ISI, during the first 2 spikes of each depolarizing phase;
- the *f-I_stim_* slope was derived from initial responses to the excitatory step currents;
- the *f_ss_/f* ratio was used to evaluate spike-frequency adaptation, where *f_ss_* was computed at the end (last 5 spikes) of the first 1-sec interval of depolarizing stimulation;
- the parameter 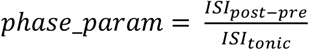 was computed to evaluate phase-reset; *ISI_post-pre_* was the interval between the 2 spikes preceding and following the impulse current, while *ISI_tonic_* was the average ISI during the 2.5-s zero-input intervals of phase-reset testing (Fig. 2D);
- latency and frequency of 1^st^ spike were measured in the rebound burst after hyperpolarization (*lat_rebound* and *rebound_freq*, respectively);
- the response speed was computed to evaluate resonance, as the inverse of the mean spike latency in each resonance step; then the values from multiple simulation tests and frequencies were fitted through a smoothing spline in order to obtain the resonance curve (Gandolfi et al., 2013).

Parameter values for simulations and recordings are reported as mean ± SD.

In addition, for quantifying experimental subthreshold oscillations, the power spectral density of 1-sec *V_m_* traces was computed and, for each recording, the main oscillation frequency was associated to the spectrum peak.

## 3 Results

The E-GLIF neuron is a linear mono-compartment neuron model using three differential equations for membrane potential (*V_m_*), a depolarizing current (*I_dep_*) and an adaptation current (*I_adapt_*). These state variables are updated at each spike event, which occurs according to a probabilistic threshold crossing controlled by escape noise. E-GLIF contains 6 fixed parameters (*C_m_, τ_m_, E_L_*, Δ*t_ref_, V_th_, V_r_*) and 6 tunable parameters (*k_adap_, k_2_, A_2_, k_1_, A_1_, I_e_*) and is designed to obtain a flexible representation of complex firing dynamics. The equation system is analytically solvable. The interdependency across state variables makes the model able to show oscillatory behaviors. The update mechanisms occurring at spike events, that are specific for each state variable, generate adaptive behaviors on multiple time scales.

In the present development, an optimization strategy using a gradient-based minimization method allowed full control over the search of optimal parameters. This operated with explicit solutions related to specific time frames making unnecessary to compute the state variables at each time instant throughout a simulation. Desired spike times were imposed depending on input stimulation.

To challenge E-GLIF in a complex and meaningful case, the cerebellar GoC was chosen as the target neuron to be modeled. Once extracting the optimal parameters, a complete simulation of the neuron could be carried out by applying a multi-phase stimulation protocol able to reveal the emergence of the various spiking response patterns. An identical protocol was used to stimulate real GoCs in acute cerebellar slices, so as to obtain experimental data for robust model validation.

### 3.1 Optimization

In order to reproduce non-damped membrane potential oscillations, *k_2_* was set to 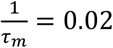 (Section 2.3.1). For the remaining parameters, multiple optimization runs with random initialization converged towards similar values for all the 5 runs within 200 iterations, without achieving values on the boundary of the search space (Fig. 3A-E). Optimization, exploring the 5-D space of tunable parameters, stably met minimization of the cost function and compliance with all the mutual parameter constraints: all the optimization runs ended with a cost function value lower than 0.5 and all the constraints verified (Fig. 3F). Considering the median of the final parameter sets, the resulting parameter values were: *k_adap_, A_2_, k_1_, A_1_, I_e_* = [0.22, 178.01, 0.03, 259.99, 16.21].

**Figure 3.**
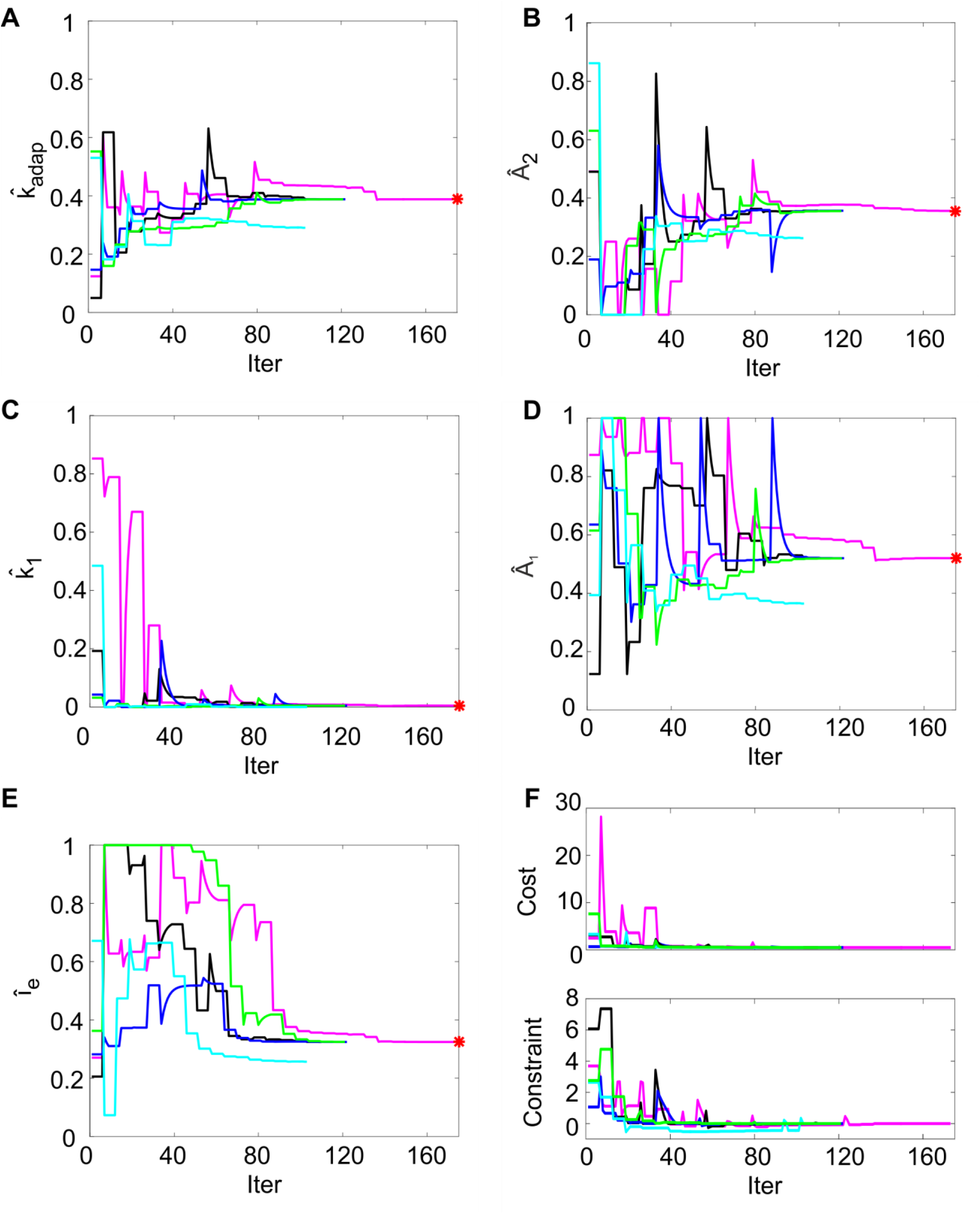
Parameter optimization. (A)-(E) Tunable parameters evolution over 5 optimization runs. Normalized parameter values (*k_adap_, A_2_, k_1_, A_1_, I_e_*) are represented along iterations. In each run (different colors), the parameter starts from random values within permitted ranges (shown normalized on y-axis). In each panel, the red point represents the optimal final parameter value (median across optimization runs). (F) Cost and constraints functions over 5 optimization runs. The cost and the constraint functions are reported for the iterations in 5 optimization runs (different colors). Especially in the starting iterations, the optimization algorithm tries to minimize the cost function while respecting the constraints. Parameter search can cause evident changes in the cost function value, especially in the first explorative iterations.

### 3.2 Model responsiveness during current step protocols

The optimized model was tested by 10 simulations in PyNEST (Eppler et al., 2009), letting *V_m_* and the two synthetic currents to evolve in time and to be updated at each spike, thus generating corresponding spike patterns, following the stochastic rule (see Section2.1).

Using the parameters resulting from the optimization process, the model was able to generate a linear relationship between *I_stim_* and response frequency *f*, with a constant slope of 0.2 Hz/pA (Fig. 4A). The autorhythm was at frequency *f_tonic_* = 12.8 ± 0.02 Hz (Fig. 4A and 4B). The escape noise process caused a slight ISI variability (CV_ISI_ = 3.4 ± 1.4% during the zero-current phase) with the parameters *λ_0_* and *τ_V_* set to 1 and 0.4, respectively.

**Figure 4.**
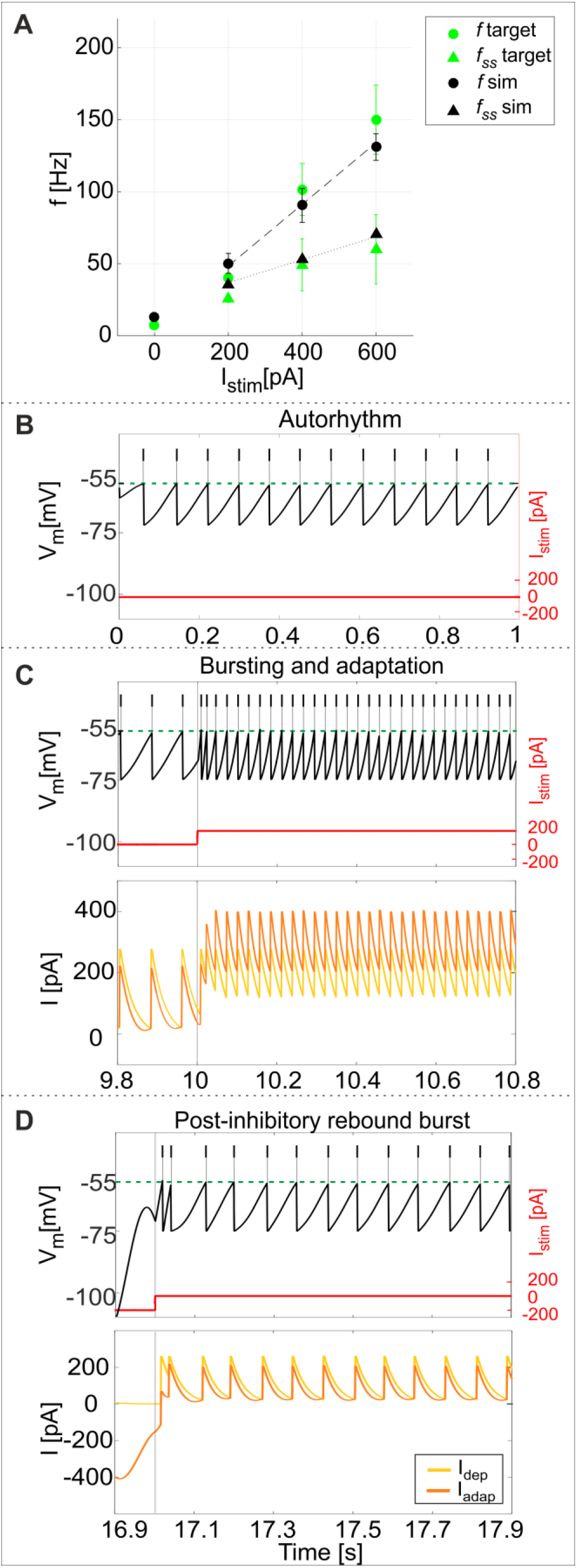
PyNEST simulations. (A) *f-I_stim_* relationships. Responses of the optimized GoC neuron during continuous simulations in PyNEST including firing stochasticity. For each stimulation current step (zero-current, 200, 400 and 600 pA lasting 1 second each), the frequency of spiking activity at the beginning (dot markers - *f*) and at the end (triangle markers – *f_ss_*) is computed. Black: simulation results (mean and SD across 10 simulation runs, with low variability in the autorhythm and steady state frequencies). Green: target values used in the optimization (mean and S<u>D</u>d of the target value distribution). The difference between instantaneous frequencies of the first spikes (*f*) and of the last spikes at 1 second (*f_ss_*) is an estimation of the spike-frequency adaptation. (B)-(D) Model responses to different stimulations. Membrane potential (black), spike events (black lines in the upper part), and the input current (red) in three blow-ups (1-sec time windows) along a continuous simulation in PyNEST of the optimized GoC neuron. (B) Autorhythm phase with zero-stimulation current; (C) Depolarization-induced bursting and spike-frequency adaptation during a current step injection of 200 pA. The currents *I_dep_* (yellow line) and *I_adap_* (orange line) are plotted along with the membrane potential: *I_dep_* is faster and contributes to bursting mechanisms, while *I_adap_* (with lower update rate) generates spike-frequency adaptation reaching its steady-state value after the first 2 spikes following depolarization have already occurred. (D) Rebound burst after a negative current step of −200 pA. Depending on *V_m_*, *I_adap_* reaches negative values during hyperpolarization (resulting in a depolarizing effect on *V_m_*), and contributes to the latency of rebound burst when the external inhibitory stimulation stops, while *I_dep_* activating after the first spike, sustains the burst.

The model response at increasing *I_stim_* lasting 1 s was (mean ± SD): *I_stim_* = 200 pA, firing rate 49 ± 6 Hz at the beginning of the current step and 36 ± 0.2 Hz at the end; *I_stim_* = 400 pA, firing rate 90 ± 10 Hz at the beginning of the current step and 53 ± 0.2 Hz at the end; *I_stim_* = 600 pA, firing rate 134 ± 8 Hz at the beginning of the current step and 68 ± 0.2 Hz at the end (Fig. 4A). As shown in Figure 4C, the higher initial frequencies reflected the depolarization-induced bursting driven by *I_dep_*, while the decrease of the firing rate along the stimulation step was mainly caused by the slightly slower *I_adap_* increase and corresponds to spike-frequency adaptation, which becomes more pronounced at higher stimuli (Fig. 4A) (Solinas et al., 2007a).

After a hyperpolarizing current step (−200 pA), the model showed a post-inhibitory rebound burst before recovering to spontaneous firing rate, with latency 30 ± 13 ms and frequency 47 ± 5 Hz (Fig. 4D). This effect reflected the different dynamics of the two currents, *I_adap_* and *I_dep_*, affecting *V_m_*: the current *I_adap_*, being coupled with *V_m_*, reached negative values during inhibitory stimulation and thus contributed to depolarize the neuron when the stimulation stopped (Par. 2.1), affecting the latency of rebound burst. After the first spike, the fast current *I_dep_* sustained burst persistence; after the first 2 spikes *I_adap_* attained a steady state balance with *I_dep_*, bringing back the activity to the autorhythm (Fig. 4D).

### 3.3 Model validation against experimental data

Experimental data from 5 GoCs showed physiological inter and intra variability in recorded voltage traces under different stimulation protocols. They exhibited autorhythm at a rate of 11.5 ± 8 Hz, increasing depolarization-induced bursting and spike-frequency adaptation in response to positive current steps, rebound bursts after negative current steps (Fig. 5A-C). These features were fully reproduced by the GoC model, as demonstrated by a validation test using more numerous and different current inputs (see Fig. 2D). The autorhythm was stably reproduced (Fig. 5D and 6A); the linearity between *I_stim_* and response *f* was maintained over multiple *I_stim_* levels (Fig. 5E and 6A), confirming the results of the first simulations with a shorter stimulation protocol (Fig. 4). The rebound burst systematically occurred after a hyperpolarization and its internal speed and latency increased, with higher absolute values of the preceding negative current steps, consistent with experimental results (Fig 5F, Table 4).

**Table 4.** Validation and prediction (e.g. phase reset with higher pulse) of rebound bursting and phase reset mechanisms.

| *Parameter* | *Protocol phase* | **Experimental**<br>**(mean±SD)** | **Simulation**<br>**(mean±SD)** |
| --- | --- | --- | --- |
| *lat_rebound [ms]* | after *inh* = -100 pA | 20.5±8.5 | 21.9±5.6 |
|  | after *inh* = -200 pA | 24.1±8.3 | 26.3±10.9 |
| *rebound_freq [Hz]* | after *inh* = -100 pA | 36.8±17.2 | 30.4±3 |
|  | after *inh* = -200 pA | 44.2±18.7 | 45.6±7.7 |
| *phase_param* | after pulse = 4000 nA | 1.65±0.38 | 1.84±0.09 |
|  | after pulse = 6800 nA | not acquired | 1.9±0.16 |

**Figure 5.**
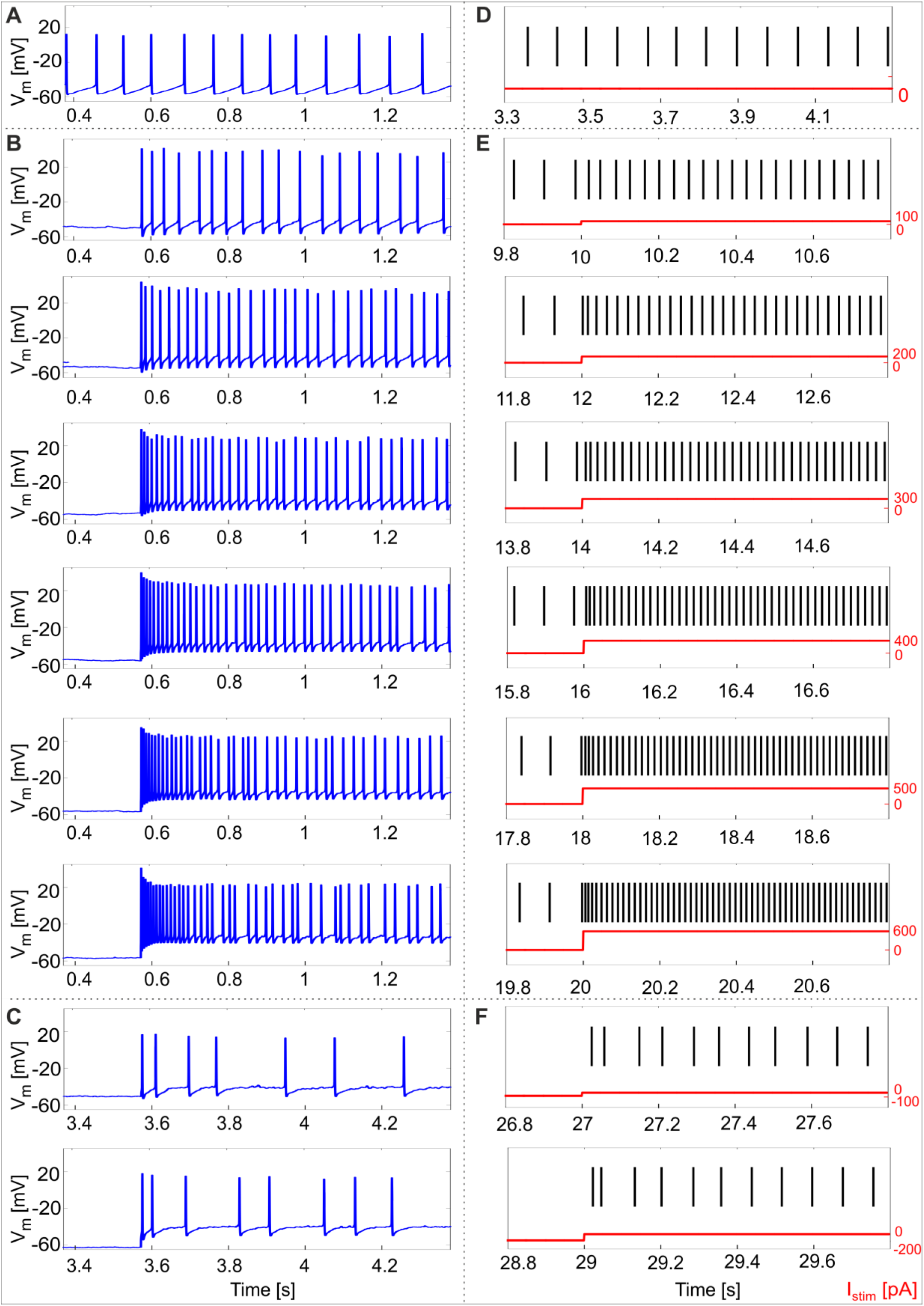
Responses to current steps. (A)-(C) Example of GoC membrane potential recorded during *in vitro* experiments under different step current stimulation protocols. (A) Autorhythm; (B) Depolarization induced bursting and spike-frequency adaptation with increasing current steps from 100 pA to 600 pA (with increments of 100 pA); (C) Rebound burst after a negative current step of −100 pA and −200 pA. (D)-(E) Simulated spike patterns during 1-sec time windows of the multi-phase stimulation protocol, with different input step currents used also in the recordings (red). (D) Autorhythm with ISIs irregularity. (E) Depolarization-induced bursting and spike-frequency adaptation at increasing levels of input current (from 100 pA to 600 pA, with increments of 100 pA). Reflecting experimental behavior, a clear firing rate increase is present as the stimulation starts, soon decreasing to the steady state value. Both initial and final spiking frequencies are higher with increasing input currents. (F) Post-inhibitory rebound bursting following a hyperpolarization of −100 pA (upper panel) and −200 pA (lower panel): bursting properties are enhanced after a stronger inhibitory current step, coherently with experimental results.

**Figure 6.**
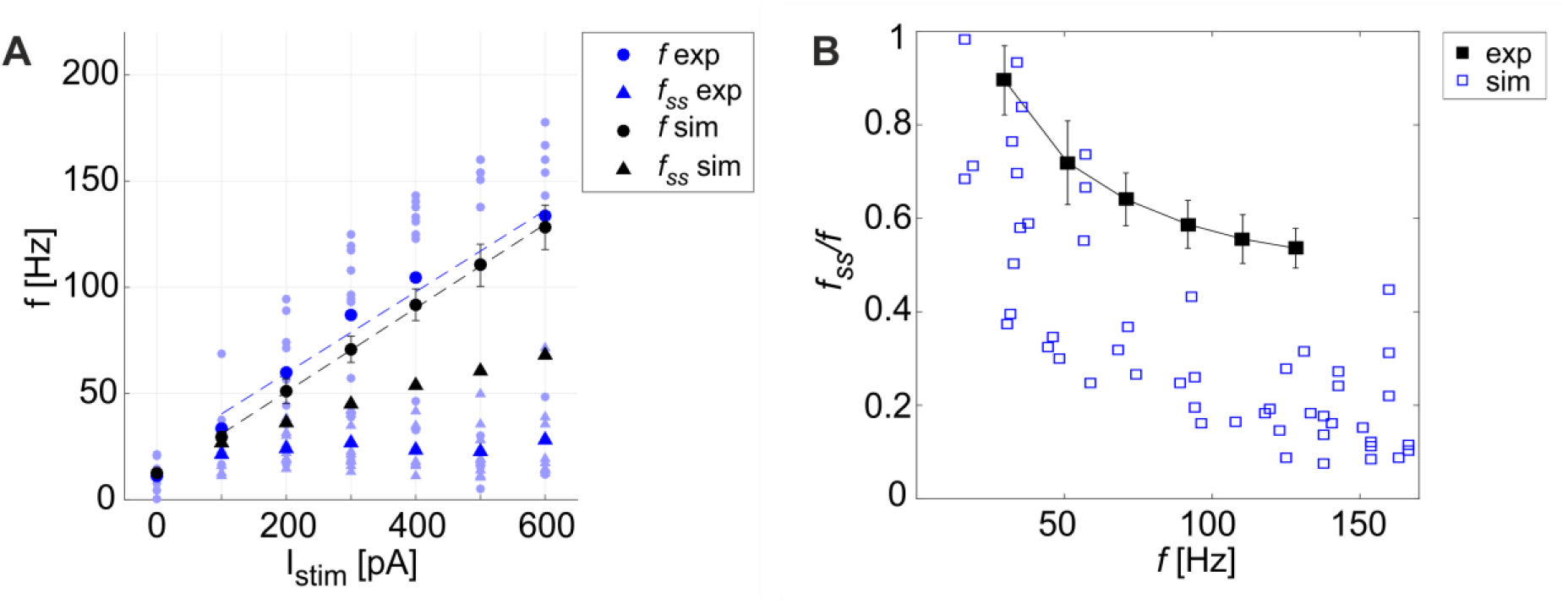
Excitability and adaptation. (A) *f-I_stim_* curve: different firing rates at the beginning (*f*) and after 1 second (*f_ss_*) of stimulation with increasing step currents, in mouse recordings (mean value highlighted in blue, all experimental data in light blue) and simulations (in black, mean and SD in 10 simulations). Linear fitting curves for both simulations and experimental data (dashed lines) result in the same *f-I_stim_* slope (0.2 Hz/pA). (B) Spike-frequency adaptation mechanisms: comparison of adaptation rate in experiments (blue) and model simulations (mean and SD of 10 simulations, in black). Both in A and B, the imperfect match between recordings and model is likely to reflect steady-state experimental degeneration not accounted for by the model.

A linear fitting on the experimental data resulted in the same *f-I_stim_* slope (0.2 Hz/pA) as when fitting simulation data (Fig. 6A). Experimental recordings and model behaviors evidently differed in the steady-state response rate (*f_ss_*) that may be related to experimental mechanisms not modelled in simulations. Indeed, the experimental *f_ss_* values were lower than in the model (Fig. 6A), thereby reducing adaptation in the model compared to experiments (Fig. 6B).

An important phenomenon evident in GoCs (as well as in some other brain neurons) is *phase-reset*, which allows desynchronization within sub-circuits triggered by strong impulses (Fig. 7A) (Buzsáki, 2004; Buzsáki and Draghun, 2004). The GoC E-GLIF was able to reproduce this feature (Fig. 7B), thanks to the coupling between *I_adap_* and *V_m_* that caused a rapid increase of *I_adap_* when *V_m_* value raised following a huge external pulse; this blocked spike generation for the same time interval independent from the phase of autorhythm before the pulse, resetting the cells phase of the autorhythm. Impulse amplitude was an element affecting the afterhyperpolarization duration following the pulse-triggered spike, slightly increasing the phase-reset parameter as the pulse charge increased (Table 4). The E-GLIF simulations matched the experimental results with the impulse at 4000 pA and predicted the slightly increased phase-reset in case of higher pulse current (6800 pA), confirming literature results (Solinas et al., 2007b).

**Figure 7.**
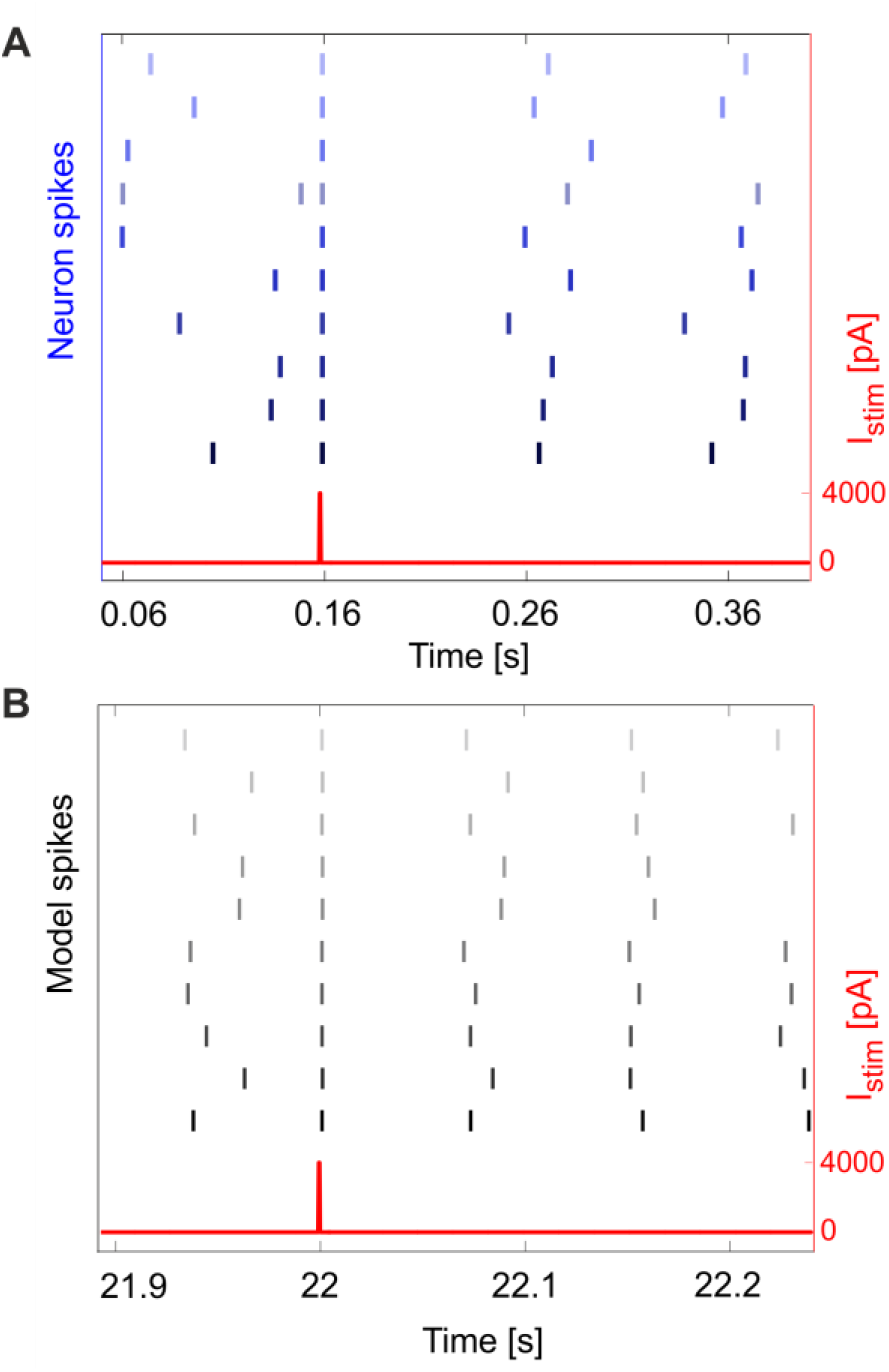
Phase reset. Phase reset in a series of 10 simulations (A) and experimental acquisitions (B), with input current pulse of amplitude *I_stim_* = 4000 pA. In panel A, we show the input current and the neuron spikes derived from 10 recorded *V_m_* traces, while in panel B we show the input current and spike events (gray lines) for 10 simulations. In both cases, at the impulse time, the neuron fires, thus realigning the spike event to the input, independently from the phase of the rhythmic firing before the impulse.

Finally, another property evident in GoCs and often observed in central nervous system neurons is the presence of endogenous *subthreshold oscillations*, which provide a fundamental mechanism for efficient network intercommunication and plasticity. Oscillations are correlated to *resonance*, and both have been shown to depend on intrinsic membrane properties (Hutcheon and Yarom, 2000). The cerebellar GoCs exhibit intrinsic theta-frequency subthreshold oscillations and resonance (Solinas et al., 2007b) that are thought to be instrumental to propagate similar properties throughout the entire cerebellum granular layer (Gandolfi et al., 2013). GoCs work as a band-pass filters by amplifying the input in the theta band. The present experimental data showed *V_m_* subthreshold oscillations at 4.2 ± 1.4 Hz (Fig. 8A): the normalized power spectral density from 9 experimental *V_m_* traces had a peak between 2 and 6 Hz (theta band). As a result, during 5 recordings with the periodic stimulation protocol, the average response speed curve (evaluating resonance) exhibited a maximum at 3.5 Hz (Fig. 8B) and all the curves in each individual recording also had the highest value at 3.5 Hz. Consistently, the simulated GoC model behaved as a band-pass filter with a prominent peak at 3.5 Hz (Fig. 8D), while exhibiting pure *V_m_* self-sustained oscillations at 5.5 Hz (Fig. 8C) (Session 2.3.1). In order to verify the low- and high-pass properties of the f_stim_-response rate curves, the model was simulated with additional stimulation frequencies and generated points falling within the predicted band-pass filter region (Fig. 8D). It should be noted that the slight difference among the frequencies of autorhythm, subthreshold oscillations and resonance (Hutcheon and Yarom, 2000) observed in the experimental data [for a similar effect see (Solinas et al., 2007a, 2007b)] was also observed in model simulations.

**Figure 8.**
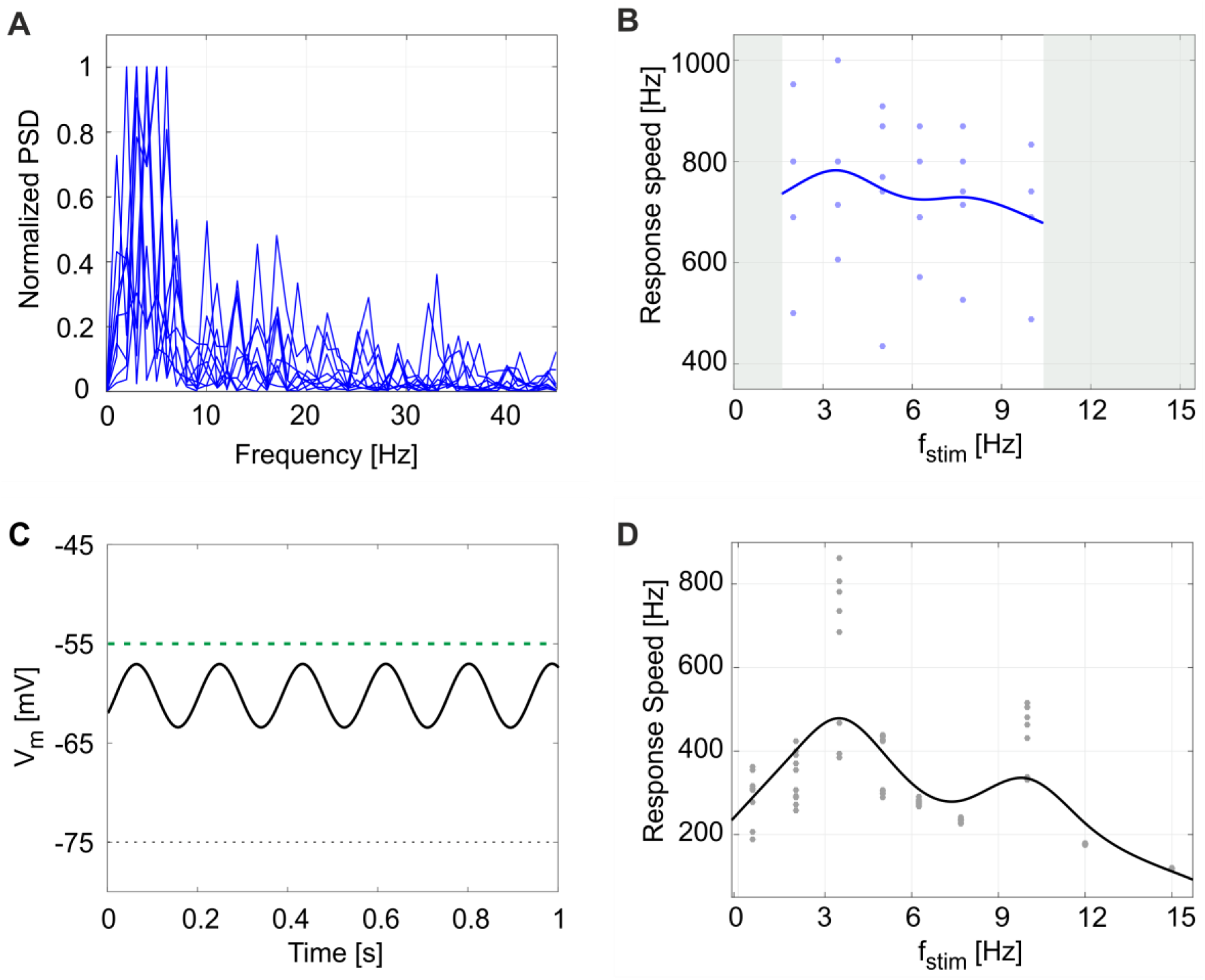
Subthreshold oscillations and resonance. (A) Normalized power spectral density (PSD) of 1-sec membrane potential traces from 9 experimental recordings. All plots show a peak in theta band (between 2 and 6 Hz). (B) Response speed values for 5 experimental recordings with the resonance protocol (using f_stim_ from 2 to 10 Hz), and fitted resonance curve showing a maximum at 3.5 Hz, despite data variability. (C) Model membrane potential self-sustained sinusoidal subthreshold oscillations in theta band, when the spiking mechanism is blocked. (D) Response speed during 10 simulations with the periodic input pattern fitted with a smoothing spline, showing resonance in theta band (higher peak at 3.5 Hz). Simulation results confirms the low- and high-pass effect in the frequency ranges not acquired during experiments (gray areas in panel B).

### 3.4 Model responsiveness to synaptic stimulation

In order to simulate synaptic responsiveness, synaptic weights and delays were set to different values for both excitatory and inhibitory synapses (excitatory synapses 40 nS and 0.1 ms, inhibitory synapses 10 nS and 0.1 ms). As shown in Fig. 9B, synaptic stimulation of E-GLIF caused an alpha-shaped change in synaptic conductance. The excitatory spike train increased irregularity in neuron firing (CV_ISI_ = 39%), consistent with the irregular spiking of GoCs *in vivo* (Cerminara and Rawson, 2004; D’Angelo, 2009). The inhibitory spike train generated a rebound burst in E-GLIF after the end of the input stimulus, due to the intrinsic changes of the model currents. This result confirmed the ability of GoC E-GLIF to reproduce the rich variety of electrophysiological properties of GoCs.

**Figure 9.**
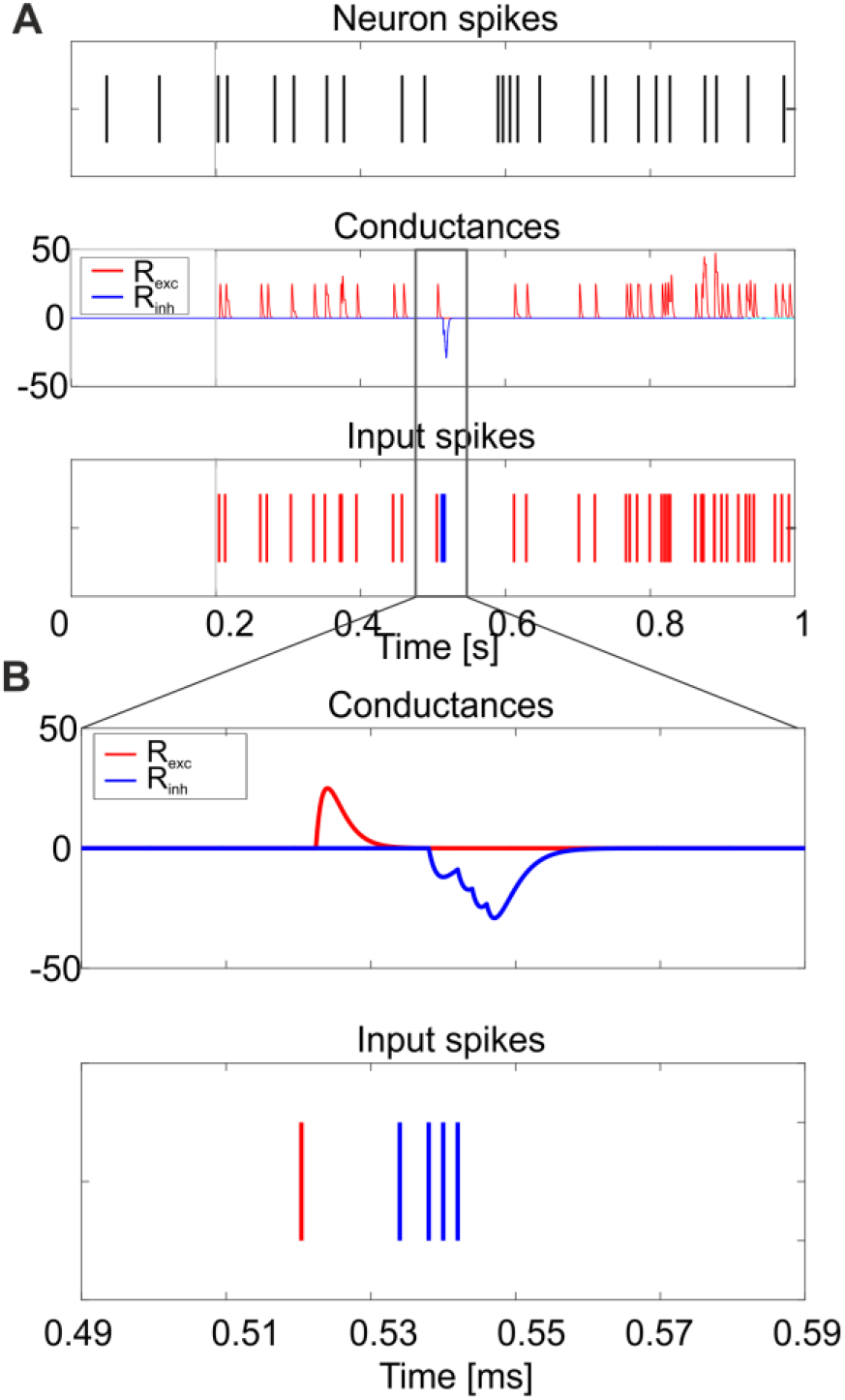
Responses to synaptic inputs. Simulation of the neuron model response to synaptic inputs on two synaptic receptor mechanism (R_exc_ and R_inh_). The synaptic models are conductance-based, with an alpha-shaped conductance change. As shown in panel (A), the excitatory spike train (bottom panel - red) increases irregularity in neuron firing (top panel), while the inhibitory input burst (bottom panel - blue) produces the rebound burst. Synaptic conductances are modified according to an alpha function, with delay 0.1 ms and maximum change depending on the weight of the synapse (middle panel). A zoom on the conductance change is shown in panel (B).

## 4 Discussion

This work reports the development, optimization and testing of the *extended generalized leaky integrate-and-fire* neuron model, E-GLIF. E-GLIF is a simplified point-neuron based on a system of 3 linear ordinary differential equations that is able to represent multiple complex electrophysiological mechanisms at different levels of abstraction. Like GLIF, E-GLIF maintains analytical tractability, allows to define different solution regimes and to optimize model parameters through minimization methods. The main improvement provided by E-GLIF is to generate a richer set of neuronal dynamics beyond depolarization-induced bursting and adaptation, which includes also rebound bursting, phase-reset, intrinsic sub-threshold oscillations and resonance. Thus, E-GLIF covers almost the whole set of neuronal discharge properties relevant for microcircuit functioning and network entrainment. Moreover, E-GLIF is designed to maintain a traceable correspondence between lumped parameters and ionic conductances of the neuronal membrane. In this way, E-GLIF allows to combine GLIF computational efficiency with the ability of reproducing the salient dynamic properties of neuronal discharge, while keeping insight into the underlying cellular mechanisms. E-GLIF appears therefore suitable to bridge the gap between biophysically detailed realistic models and computationally efficient simplified models, and could be used to investigate the impact of neuronal dynamics in large-scale networks. As a prove of its validity, E-GLIF was shown to reproduce the main spiking discharge properties of the cerebellar GoC, a prototype of a neuron with complex electroresponsiveness.

### 4.1 Model parameterization and optimization strategy

By exploiting its analytical solution, the E-GLIF GoC model was optimized using gradient descent minimization methods. This allowed to fast-tune a unique set of parameters generating the appropriate spiking responses to various input patterns. The cost function was designed to evaluate sub-threshold membrane potential dynamics using the integral of membrane potential all over the inter-spike interval: in this way, the cost function was still differentiable, while taking into account the full history of the signal preceding the spike. Supra-threshold dynamics may not be relevant since E-GLIF used a spike-update-reset mechanism. In addition, the general cost function could be customized by disabling the terms corresponding to the properties that are not exhibited by the specific neuron model. For example, for a neuron without autorhythm, the *error_zero_stim_* term would not be included in the cost function. For the remaining terms, the desired output parameters listed in Table 1 were derived from electrophysiological studies reported in literature and generally applicable to many different neuron types. It should also be noted that, although stochasticity in spike generation was not expressed in the explicit model solution used for optimization, it was accounted for by optimizing the neuron on a distribution of desired spike times.

Compared to other optimization strategies, which are based on semi-automatic fitting on a training set of neural traces or spiking patterns (Pozzorini et al., 2015; Rössert et al., 2016) or realistic model mapping (Marasco et al., 2012), E-GLIF tuning was based on features related to neurophysiological activity (primarily membrane capacitance and time constant, resting potential, refractory period, spike threshold) and on input-output relationships. Then, the model was validated against real experimental data. This *feature-based* approach guarantees high generalization capability and would be particularly suitable when the data needed for SNN models reconstructions have been obtained in different laboratories and therefore do not share the uniform database required for automatic fitting and validation.

### 4.2 Comparison to other simplified models

E-GLIF integrates elements taken from the theory of point neuron models in order to reach a good compromise between computational efficiency, number of tunable parameters and biological plausibility (Izhikevich, 2004; Pozzorini et al., 2015; Teeter et al., 2018).

First, E-GLIF reproduces sub-threshold responses and spike timing rather than the shape of action potentials. Nonetheless, some spike properties, such as spike width and afterhyperpolarization, are accounted for by the reset rules, which map the state after the spike to that before it (Teeter et al., 2018). It should be noted that inclusion of adaptive exponential terms, useful to reproduce supra-threshold dynamics (Brette and Gerstner, 2005), is not needed in E-GLIF, which is conceived for embedding into SNN and therefore privileges spike timing and population spiking patterns.

Secondly, the E-GLIF analytical tractability can be exploited to implement versions designed for time-driven simulations, where the exact spike time can be computed through iterative methods like bisection (Hanuschkin et al., 2010). This would be useful to decrease simulation time resolution without losing spike timing accuracy in large-scale SNNs.

Thirdly, E-GLIF exploits an adaptive current coupled with membrane potential to model spiking responses driven by adaptation mechanisms, like spike-frequency adaptation and post-inhibitory rebound bursting, rather than a sliding threshold depending on the actual membrane voltage (Mihalaş and Niebur, 2009; Pozzorini et al., 2015). This E-GLIF feature, in addition to decreasing the number of ODEs in the model, correlates effectively to the membrane mechanisms of firing, as exemplified in the case of cerebellar GoCs (cf. (Solinas et al., 2007a, 2007b)). In GoCs, both the the *f-I_stim_* curve shift during adaptation and post-inhibitory rebound bursting are driven by the adapting current *I_d<u>ad</u>apt_*, in agreement with ionic mechanisms revealed by data-driven realistic modeling (Solinas et al., 2007a) and with mechanisms adopted in other simplified models (Benda et al., 2010; Naud and Gerstner, 2012)(Naud and Gerstner, 2012; Solinas et al., 2007a)(Naud and Gerstner, 2012; Solinas et al., 2007a)(Naud and Gerstner, 2012; Solinas et al., 2007a). Moreover, the adaptive current coupled with membrane potential allowed to simulate intrinsic self-sustained subthreshold oscillations at a preferred frequency and thus to generate intrinsic resonance. Importantly, this is not possible either in the traditional GLIF neurons (Pozzorini et al., 2015) or in other nonlinear adaptive point neurons (Touboul, 2012). The spike-triggered updates of *I_adap_* and *I_dep_* can be associated to mechanisms of ion-channel activation/deactivation, in particular concerning K^+^ (slow) and Na^+^ (fast) currents (Mihalaş and Niebur, 2009), as further considered below.

Finally, for testing the neuron with synaptic inputs, conductance-based synaptic receptors were embedded into the model. It should be noted that current-based synapses may be used to further improve computational efficiency at the expense of biophysical realism (Cavallari et al., 2014).

### 4.3 Correspondence of model parameters to subcellular mechanisms

Given the specific design of E-GLIF (see above), it is possible to trace the relationship between E-GLIF parameters and the ionic and membrane mechanisms generating the spiking response in real neurons and realistic models. This comparison is well exemplified by considering the ionic mechanisms of cerebellar GoCs, which have been previously determined in great detail (Solinas et al., 2007a, 2007b). Table 5 shows a list of membrane currents generating various electrophysiological properties in the cerebellar GoC (D’Angelo et al., 2013) and their remapping onto the lumped parameters of E-GLIF, at a different level of abstraction.

**Table 5.** Correspondence between subcellular mechanisms that generated specific electrophysiological properties and how they were simplified in E-GLIF model elements. The arrow indicate depolarizing (→) and hyperpolarizing (↓) actions of the membrane currents in the real cell and models. I_Na-t_: transient sodium current; I_K-V_: delayed rectifier potassium current; I_h_: hyperpolarization-activated current; I_Na-p_: persistent sodium current; I_K-slow_: slow M-like potassium current; I_Ca-HVA_: high voltage-activated calcium current; I_K-AHP_: SK-type calcium-dependent potassium current; I_Na-r_: resurgent sodium current; I_K-A_: A-type potassium current; definitions and properties of the ionic currents are given in (Forti et al., 2006; Solinas et al., 2007a, 2007b).

| **Ionic channel mechanisms** | **Electrophysiological property** | **E-GLIF mechanisms** |
| --- | --- | --- |
| $I_{Na-t} \uparrow / I_{K-v} \downarrow$ balance | Action potential | Digital |
| $I_h \uparrow$<br>$I_{Na-p} \uparrow / I_{K-slow} \downarrow$ balance<br>$I_{Ca-HVA} \uparrow / I_{K-AHP} \downarrow$ balance | Autorhythmicity | $I_e \uparrow / I_{adap} \downarrow$ balance |
| $I_{Na-r} \uparrow$ and $I_{K-A} \downarrow$ | Depolarization-induced burst | $I_{dep} \uparrow$ |
| $I_{Ca-HVA} \uparrow / I_{K-AHP} \downarrow$ balance<br>$I_{K-slow} \downarrow$ | Spike-frequency adaptation | $I_{adap} \downarrow$ |
| $I_{Ca-HVA} \uparrow / I_{K-AHP} \downarrow$ balance | Phase-reset | $I_{adap} \downarrow$ |
| $I_h \uparrow$<br>$I_{Ca-LVA} \uparrow$ | Post-inhibitory rebound burst | $I_{adap} \uparrow$ and $I_{dep} \uparrow$ |
| $I_{K-slow} \downarrow$ and $I_{Na-p} \uparrow$ | Subthreshold oscillations | $V_m - I_{adap}$ coupling |
| $I_{K-slow} \downarrow$ and $I_{Na-p} \uparrow$ | Resonance | $V_m - I_{adap}$ coupling |

While, in order to simultaneously generate 8 different electrophysiological properties (action potential, autorhythmicity, depolarization-induced burst, post-inhibitory rebound burst, spike-frequency adaptation, phase-reset, subthreshold oscillations, resonance), the realistic GoC model engages 10 ionic currents (I_h_, I_Na-t_, I_Na-p_, I_Na-r_, I_Ca-HVA_, I_Ca-LVA_, I_K-V_, I_K-A_, I_K-AHP_, I_K-slow_), E-GLIF has just 3 currents (*I_e_, I_adap_, I_dep_*) controlling a digital spike generator. Actually, there are correlations between ionic mechanisms (e.g. I_Ca-HVA_ and I_K-AHP_ are coupled), the same ionic mechanisms can influence multiple electrophysiological properties (e.g. the I_Ca-HVA_ /I_K-AHP_ balance influences both autorhythmicity, adaptation and phase-reset), and some electroresponsive properties are at least partly bound one to each other (e.g. subthreshold oscillations and resonance) reducing the dimensionality of the real neuron parameter space (Solinas et al., 2007a, 2007b). *De facto*, with its 3 currents and spike-reset mechanisms, E-GLIF can effectively abstract the high-dimensional response pattern of the GoC, supporting the concept that appropriate models can provide a mathematical interpretation of complex neuronal properties (Gerstner and Naud, 2009). It should be noted that other simplified non-linear models (e.g. (Izhikevich, 2003)), appear to be less interpretable in mechanistic terms and less flexible in generating complex response patterns. In addition, they associate regions of the parameter space to different spiking regimes, instead of generating different firing responses based only on the input stimulus.

### 4.4 Conclusions

This work shows that it is possible to represent complex neuronal firing dynamics through a mono-compartment neuron model by updating the GLIF into E-GLIF at the modest computational expense of 3 ODEs. Yet there is a remarkable efficiency gain of 10^2^-10^3^ times with respect to realistic multi-compartmental models. For example, compared to the realistic version of the GoC model (Solinas et al., 2007a, 2007b) with 23 ODEs, there is an 80-fold computational time reduction to simulate the same stimulation protocol with the E-GLIF. Together with the computational efficiency, the E-GLIF was able to reproduce multiple electrophysiological features with a single set of model parameters, moving forward the traditional approach of neuron modelling and resulting in a higher biological plausibility (Izhikevich, 2004). Specific advantages of E-GLIF are the second-order dynamics and the linearity: the model admits an oscillatory response to constant inputs and an analytical solution that allows to extend the theoretical analysis of complex firing dynamics. Moreover, E-GLIF keeps a correspondence between lumped model parameters and electrophysiological mechanisms, limiting black-box fitting and supporting the interpretation of neuronal physiological properties and their changes by neuromodulation and plasticity. There is a large category of neurons that could be represented as point processes and could indeed be modeled with E-GLIF. For example, good candidates are the granule cells and the stellate and basket cells in the cerebellum as well as various interneurons in the neocortex and hippocampus. Nonetheless, when dendritic or axonal computations become critical, E-GLIF modifications would be needed, e.g. by connecting multiple E-GLIF style compartments one to each other and adopting dendritic compression and synaptic efficacy scaling procedures (Marasco et al., 2012). This could be the case of Purkinje cells (Masoli et al., 2015; Masoli and D’Angelo, 2017) in the cerebellum and of pyramidal cells in the neocortex and hippocampus (Migliore et al., 2008). Together with the other GLIF neurons, E-GLIF could contribute to generate a standardized database of computationally efficient models capable of generating a rich repertoire of non-linear firing patterns applicable to diverse brain regions and scientific issues by the community (Pozzorini et al., 2015; Rössert et al., 2016; Teeter et al., 2018; Tiesinga et al., 2015). Along this line, the implementation of E-GLIF on the NEST platform (Diesmann and Gewaltig, 2002) is expected to bring salient aspects of neuronal firing dynamics into large-scale network simulations, where different point neuron models can be combined based on the complexity of the neural population to be represented (Destexhe et al., 1996; Geminiani et al., 2017). The resulting SNN models could help understanding how brain networks generate their computations. Indeed, E-GLIF has been designed to investigate cerebellar network dynamics during closed-loop behavioral testing in neurorobots (Antonietti et al., 2016; Casellato et al., 2014, 2015; D’Angelo et al., 2016b). In conclusion, E-GLIF could effectively bridge bottom-up and top-down modeling approaches (Herz et al., 2006) paving the way to the establishment of a set of simplified yet biologically meaningful spiking neuron models as the fundamental elements of multi-scale brain modeling.

## Author contribution

AG and CC elaborated the mathematical model and optimization, carried out the simulations analyzing them together with the experimental dataset, and substantially contributed to text writing. FL and FP carried out experimental recordings and analysis. AP coordinated mathematical modelling and simulations and substantially contributed to the writing of the final manuscript. ED coordinated the whole work and wrote the final version of the manuscript.

## Conflict of Interest

The authors declare that the research was conducted in the absence of any commercial or financial relationships that could be construed as a potential conflict of interest.

## Acknowledgements

This project has been developed within the *CerebNEST* HBP Partnering Project and has received funding from the European Union’s Horizon 2020 Framework Programme for Research and Innovation under Grant Agreement No. 720270 and 785907 (Human Brain Project SGA1 and SGA2).

## References

Antonietti, A., Casellato, C., Garrido, J. A., Luque, N. R., Naveros, F., Ros, E., et al. (2016). Spiking Neural Network with distributed plasticity reproduces cerebellar learning in Eye Blink Conditioning paradigms. IEEE Trans. Biomed. Eng. 63, 210–9. doi:10.1109/TBME.2015.2485301.

Benda, J., Maler, L., and Longtin, A. (2010). Linear Versus Nonlinear Signal Transmission in Neuron Models With Adaptation Currents or Dynamic Thresholds. J. Neurophysiol. 104, 2806–2820. doi:10.1152/jn.00240.2010.

Brette, R., and Gerstner, W. (2005). Adaptive Exponential Integrate-and-Fire Model as an Effective Description of Neuronal Activity. J. Neurophysiol. 94, 3637–3642. doi:10.1152/jn.00686.2005.

Burkitt, A. N. (2006). A review of the integrate-and-fire neuron model: I. Homogeneous synaptic input. Biol. Cybern. 95, 1–19. doi:10.1007/s00422-006-0068-6.

Buzsáki, G. (2004). Large-scale recording of neuronal ensembles. Nat. Neurosci. 7, 446–451. doi:10.1038/nn1233.

Buzsáki, G., and Draghun, A. (2004). Neuronal Oscillations in Cortical Networks. Science (80-.). 304, 1926–1930. doi:10.1126/science.1099745.

Casellato, C., Antonietti, A., Garrido, J. A., Carrillo, R. R., Luque, N. R., Ros, E., et al. (2014). Adaptive robotic control driven by a versatile spiking cerebellar network. PLoS One 9, 1–17. doi:10.1371/journal.pone.0112265.

Casellato, C., Antonietti, A., Garrido, J. A., Ferrigno, G., D’Angelo, E., and Pedrocchi, A. (2015). Distributed cerebellar plasticity implements generalized multiple-scale memory components in real-robot sensorimotor tasks. Front. Comput. Neurosci. 9, 1–14. doi:10.3389/fncom.2015.00024.

Cavallari, S., Panzeri, S., and Mazzoni, A. (2014). Comparison of the dynamics of neural interactions between current-based and conductance-based integrate-and-fire recurrent networks. Front. Neural Circuits 8, 1–23. doi:10.3389/fncir.2014.00012.

Cerminara, N. L., and Rawson, J. A. (2004). Evidence that Climbing Fibers Control an Intrinsic Spike Generator in Cerebellar Purkinje Cells. J. Neurosci. 24, 4510–4517. doi:10.1523/JNEUROSCI.4530-03.2004.

Cesana, E., Pietrajtis, K., Bidoret, C., Isope, P., D’Angelo, E., Dieudonne, S., et al. (2013). Granule Cell Ascending Axon Excitatory Synapses onto Golgi Cells Implement a Potent Feedback Circuit in the Cerebellar Granular Layer. J. Neurosci. 33, 12430–12446. doi:10.1523/JNEUROSCI.4897-11.2013.

D’Angelo, E. (2009). The critical role of Golgi cells in regulating spatio-temporal integration and plasticity at the cerebellum input stage. Front. Neurosci. 2, 35–46. doi:10.3389/neuro.01.008.2008.

D’Angelo, E., Antonietti, A., Casali, S., Casellato, C., Garrido, J. A., Luque, N. R., et al. (2016a). Modeling the cerebellar microcircuit: New strategies for a long-standing issue. Front. Cell. Neurosci. 10, 1–29. doi:10.3389/fncel.2016.00176.

D’Angelo, E., and Gandini Wheeler-Kingshott, C. (2017). Modelling the brain: Elementary components to explain ensemble functions. Riv. del nuovo Cim. 40, 273–333. doi:10.1393/ncr/i2017-10137-5.

D’Angelo, E., Mapelli, L., Casellato, C., Garrido, J. A., Luque, N., Monaco, J., et al. (2016b). Distributed Circuit Plasticity: New Clues for the Cerebellar Mechanisms of Learning. The Cerebellum 15, 139–151. doi:10.1007/s12311-015-0711-7.

D’Angelo, E., Solinas, S., Mapelli, J., Gandolfi, D., Mapelli, L., and Prestori, F. (2013). The cerebellar Golgi cell and spatiotemporal organization of granular layer activity. Front. Neural Circuits 7, 1–21. doi:10.3389/fncir.2013.00093.

Destexhe, A., Bal, T., McCormick, D. A., and Sejnowski, T. J. (1996). Ionic mechanisms underlying synchronized oscillations and propagating waves in a model of ferret thalamic slices. J. Neurophysiol. 76.

Diesmann, M., and Gewaltig, M.-O. (2002). NEST: An environment for neural systems simulations. T. Plesser V. Macho (Eds.), Forsch. und wisschenschaftliches Rechn. Beitrage zum Heinz-billing-pr. 2001, Vol. 58 GWDGBericht, 43–70. Available at: http://www.scholarpedia.org/article/NEST_(NEural_Simulation_Tool).

Eppler, J. M., Helias, M., Muller, E., Diesmann, M., Gewaltig, M., and Stewart, T. C. (2009). PyNEST?: A convenient interface to the NEST simulator. 2, 1–12. doi:10.3389/neuro.11.012.2008.

Forti, L., Cesana, E., Mapelli, J., and Angelo, E. D. (2006). Ionic mechanisms of autorhythmic firing in rat cerebellar Golgi cells. J. Physiol. 3, 711–729. doi:10.1113/jphysiol.2006.110858.

Gandolfi, D., Lombardo, P., Mapelli, J., Solinas, S., and D’Angelo, E. (2013). Theta-Frequency Resonance at the Cerebellum Input Stage Improves Spike Timing on the Millisecond Time-Scale. Front. Neural Circuits 7, 1–16. doi:10.3389/fncir.2013.00064.

Geminiani, A., Casellato, C., Antonietti, A., D’Angelo, E., and Pedrocchi, A. (2017). A multiple-plasticity Spiking Neural Network embedded in a closed-loop control system to model cerebellar pathologies. Int. J. Neural Syst. 27. doi:10.1142/S0129065717500174.

Gerstner, W., and Kistler, W. M. (2002). Spiking Neuron Models. Cambridge Univ. Press doi:10.1017/CBO9780511815706.

Gerstner, W., Kistler, W. M., Naud, R., and Paninski, L. (2014). Neuronal Dynamics. Cambridge Univ. Press doi:10.1017/CBO9781107447615.

Gerstner, W., and Naud, R. (2009). How good are neuron models? Science (80-.). 326, 379–380. doi:10.1126/science.1181936.

Hanuschkin, A., Kunkel, S., Helias, M., Morrison, A., and Diesmann, M. (2010). A General and Efficient Method for Incorporating Precise Spike Times in Globally Time-Driven Simulations. Front. Neuroinform. 4, 1–19. doi:10.3389/fninf.2010.00113.

Hertäg, L., Hass, J., Golovko, T., and Durstewitz, D. (2012). An approximation to the adaptive exponential integrate-and-fire neuron model allows fast and predictive fitting to physiological data. 6, 1–22. doi:10.3389/fncom.2012.00062.

Herz, A. V. M., Gollisch, T., Machens, C. K., and Jaeger, D. (2006). Modeling Single-Neuron Dynamics and Computations: A Balance of Detail and Abstraction Motion detection. Science (80-.). 314, 80–85.

Hutcheon, B., and Yarom, Y. (2000). Resonance, oscillation and the intrinsic frequency preferences of neurons. Trends Neurosci. 23, 216–222.

Izhikevich, E. M. (2003). Simple model of spiking neurons. IEEE Trans. Neural Networks 14, 1569–1572. doi:10.1109/TNN.2003.820440.

Izhikevich, E. M. (2004). Which model to use for cortical spiking neurons? IEEE Trans. Neural Networks 15, 1063–1070. doi:10.1109/TNN.2004.832719.

Jolivet, R., Rauch, A., Lüscher, H. R., and Gerstner, W. (2006). Predicting spike timing of neocortical pyramidal neurons by simple threshold models. J. Comput. Neurosci. 21, 35–49. doi:10.1007/s10827-006-7074-5.

Jordan, J., Ippen, T., Helias, M., Kitayama, I., Sato, M., Igarashi, J., et al. (2018). Extremely Scalable Spiking Neuronal Network Simulation Code: From Laptops to Exascale Computers. Front. Neuroinform. 12. doi:10.3389/fninf.2018.00002.

Lennon, W., Hecht-nielsen, R., and Yamazaki, T. (2014). A spiking network model of cerebellar Purkinje cells and molecular layer interneurons exhibiting irregular firing. Front. Comput. Neurosci. 8, 1–10. doi:10.3389/fncom.2014.00157.

Marasco, A., Limongiello, A., and Migliore, M. (2012). Fast and accurate low-dimensional reduction of biophysically detailed neuron models. Sci. Rep. 2, 1–7. doi:10.1038/srep00928.

Markram, H. (2013). Seven challenges for neuroscience. Funct. Neurol. 28, 145–151. doi:10.11138/FNeur/2013.28.3.145.

Markram, H., Muller, E., Ramaswamy, S., Reimann, M. W., Abdellah, M., Sanchez, C. A., et al. (2015). Reconstruction and Simulation of Neocortical Microcircuitry. Cell 163, 456–492. doi:10.1016/j.cell.2015.09.029.

Masoli, S., and D’Angelo, E. (2017). Synaptic Activation of a Detailed Purkinje Cell Model Predicts Voltage-Dependent Control of Burst-Pause Responses in Active Dendrites. Front. Cell. Neurosci. 11, 1–18. doi:10.3389/fncel.2017.00278.

Masoli, S., Solinas, S., and Angelo, E. D. (2015). Action potential processing in a detailed Purkinje cell model reveals a critical role for axonal compartmentalization. Front. Cell. Neurosci. 9, 1–22. doi:10.3389/fncel.2015.00047.

Migliore, M., Novara, G., and Tegolo, D. (2008). Single neuron binding properties and the magical number 7. Hippocampus 18.

Mihalaş, Ş., and Niebur, E. (2009). A Generalized Linear Integrate-and-Fire Neural Model Produces Diverse Spiking Behaviors. Neural Comput. 21, 704–718. doi:10.1162/neco.2008.12-07-680.

Naud, R., and Gerstner, W. (2012). “The Performance (and Limits) of Simple Neuron Models: Generalizations of the Leaky Integrate-and-Fire Model,” in Computational Systems Neurobiology (Springer Netherlands).

Plotnikov, D., Rumpe, B., Blundell, I., Ippen, T., Martin, J., and Morrison, A. (2016). NESTML: a modeling language for spiking neurons. arXiv Prepr. arXiv, 93–108.

Pozzorini, C., Mensi, S., Hagens, O., Naud, R., Koch, C., and Gerstner, W. (2015). Automated High-Throughput Characterization of Single Neurons by Means of Simplified Spiking Models. PLoS Comput. Biol. 11, 1–29. doi:10.1371/journal.pcbi.1004275.

Rössert, C., Pozzorini, C., Chindemi, G., Davison, A. P., Eroe, C., King, J., et al. (2016). Automated point-neuron simplification of data-driven microcircuit models. Available at: http://arxiv.org/abs/1604.00087.

Roth, A., and van Rossum, M. C. W. (2013). “Modeling Synapses,” in Computational Modeling Methods for Neuroscientists, ed. E. De Schutter (MIT Press).

Solinas, S., Forti, L., Cesana, E., Mapelli, J., Schutter, E. De, and Angelo, E. D. (2007a). Computational reconstruction of pacemaking and intrinsic electroresponsiveness in cerebellar golgi cells. Front. Cell. Neurosci. 1. doi:10.3389/ne.

Solinas, S., Forti, L., Cesana, E., Mapelli, J., De Schutter, E., and D’Angelo, E. (2007b). Fast-reset of pacemaking and theta-frequency resonance patterns in cerebellar golgi cells: simulations of their impact in vivo. Front. Cell. Neurosci. 1, 4. doi:10.3389/neuro.03.004.2007.

Teeter, C., Iyer, R., Menon, V., Gouwens, N., Feng, D., Berg, J., et al. (2018). Generalized leaky integrate-and-fire models classify multiple neuron types. Nat. Commun. 9, 709. doi:10.1038/s41467-017-02717-4.

Tiesinga, P., Bakker, R., Hill, S., and Bjaalie, J. G. (2015). Feeding the human brain model. Curr. Opin. Neurobiol. 32, 107–114. doi:10.1016/j.conb.2015.02.003.

Touboul, J. (2012). Bifurcation analysis of a general class of nonlinear integrate-and-fire neurons. SIAM J. Appl. Math. 44. doi:10.1137/090750688.

Tripathy, S. J., Savitskaya, J., Burton, S. D., Urban, N. N., and Gerkin, R. C. (2014). NeuroElectro: a window to the world’s neuron electrophysiology data. Front. Neuroinform. 8, 1–11. doi:10.3389/fninf.2014.00040.

Zeilhofer, H. U., Studler, B., Arabadzisz, D., Schweizer, C., Ahmadi, S., Layh, B., et al. (2005). Glycinergic neurons expressing enhanced green fluorescent protein in bacterial artificial chromosome transgenic mice. J. Comp. Neurol. 482, 123–141. doi:10.1002/cne.20349.

